# The classification of orphans is improved by combining searches in both proteomes and genomes

**DOI:** 10.1101/185983

**Authors:** Walter Basile, Marco Salvatore, Arne Elofsson

## Abstract

The identification of *de novo* created genes is important as it provides a glimpse on the evolutionary processes of gene creation. Potential *de novo* created genes are identified by selecting genes that have no homologs outside a particular species, but for an accurate detection this identification needs to be correct.

Genes without any homologs are often referred to as orphans; in addition to *de novo* created ones, fast evolving genes or genes lost in all related genomes might also be classified as orphans. The identification of orphans is dependent on: (i) a method to detect homologs and (ii) a database including genes from related genomes.

Here, we set out to investigate how the detection of orphans is influenced by these two factors. Using *Saccharomyces cerevisiae* we identify that best strategy is to use a combination of searching annotated proteins and a six-frame translation of all ORFs from closely related genomes. Using this strategy we obtain a set of 54 orphans in *Drosophila melanogaster* and 38 in *Drosophila pseudoobscura*, significantly less than what is reported in some earlier studies.

## Introduction

Even after twenty years of genome sequencing some protein-coding genes appear to have no homologs. These genes are often referred to as *orphans*, a definition that dates back to the sequencing of *Saccharomyces cerevisiae* [1–3]. It was believed that most of them would disappear when more genomes would be sequenced. However that is not the case, even after thousands of genomes have been sequenced [4–7]. After the initial studies in yeast, orphans have also been identified in *Drosophila* [8–10], mammals [11], primates [12–16], plants [17, 18], Bacteria [19–21] and viruses [22, 23].

The term “orphan” might refer to genes without any detectable homologs or it might refer to only *de novo* created genes. The first definition would include genes that have homologs that all are too distant to be detected, possibly due to extensive gene loss in closely related species, lack of closely related species in the database searched, or because of fast evolution of the gene. Given better methods to detect homology and the inclusion of more closely related species in the database, the number of non-*de novo* created orphans should decrease.

If a gene truly does not have any homologs it must have been created by some mutations turning a non-coding DNA region into a protein coding gene. It is likely that this is a widespread phenomenon, at least in yeast [24]. However, in evolutionary terms only a few of these genes will get fixed in the population. It is plausible that every protein superfamily was created by such mechanisms. However, it is also possible that at least some superfamilies are created by rearranging shorter stretches of coding regions [25]. A correct identification of orphans is therefore important for understanding the selective pressures affecting novel gene creation throughout evolutionary history. In particular, an imprecise detection, that includes fast evolving genes in the orphan set, might cause the incorrect picture that *de novo* created genes have the same properties as fast evolving genes.

In some studies, different levels of orphanicity have been used [4, 26–28]. Here, a gene is assigned to have a certain age - given that homologs are only found within genomes belonging to a certain clade. For instance, a gene can be specific to primates, to mammals or eukaryotes. The correct classification of the age of a gene is difficult, as it would require accurate detection of very distant homologs. If this is not done correctly, fast-evolving genes will appear to be younger than they are. Taken these difficulties into account we only study the most recent orphans and have not used gene age in this study.

In short, the identification of orphans starts by searching for homologous genes in a set of related genomes and then labeling as orphans the genes lacking hits. The results will be dependent on the database used and the homology detection method. In general, better homology detection [29–31] would potentially reduce the number of genes assigned as orphans [4].

Another complication when it comes to identification of *de novo* created genes is that it is almost impossible to know if an ORF is a functional gene from its genomic sequence alone. For genes with homologs conservation across species is the best method to identify if a gene is functionally important or not, but by definition this is not possible for orphans. An ORF might look like a gene but is never expressed under any circumstances, i.e. not being functional. Alternatively, an ORF can be missed as it appears not to be coding. This difficulty applies both to the ORFs in the query species and to the genomes searched. Therefore, just having a hit to a genomic location does not guarantee that the corresponding region is an actual functional gene [5, 28, 32]. However, the reverse is also possible, i.e. that a hit to a non-functional ORF represents homology with a common functional ancestral gene, i.e. the function was lost in the other species but present in the ancestral one. Other factors, such as the removal of CDS overlapping annotated genes, the study of the syntenic regions, or experimental studies, improve the efficacy of detection of *de novo* created genes.

Several different estimates of the orphans of *S. cerevisiae* have been proposed [4, 5, 27, 28, 33]. Surprisingly, the obtained set of orphans is quite different between these studies. One of the reasons of these differences is the use of different datasets [28]. For example all *S. cerevisiae* ORFs are considered in Carvunis et al. [5], but only the annotated ones are present in Ekman et al. [4].

The goal of this study was to develop a robust strategy for orphan detection in sequenced genomes. We used a set of homology search methods (blastn, blastp and tblastn) and compared the classified orphans with earlier sets. To avoid some of the problems described above we decided to: (i) use only annotated genes, (ii) assume that every significant hit in a related genome shows that a gene is not *de novo* created, although we know that this would not identify very recently *de novo* created genes, (iii) use a similar set of genomes as in earlier studies. We then explore how the inclusion of different genomes and the use of different homology detection tools influence the detection of orphans first in *S. cerevisiae* and thereafter in *D. melanogaster* and *D. pseudoobscura*. At the end we propose a strategy that combines the use of tblastn and blastp. Using this strategy a conservative estimate of the number of in *Drosophila melanogaster* and *Drosophila pseudoobscura* is obtained.

## Materials and Methods

### S. cerevisiae data

5,917 *Saccharomyces cerevisiae* protein-coding genes (reference strain s288c, NCBI taxon id 559292) were downloaded from the Saccharomyces Genome Database (SGD) [34]. Only coding and annotated ORFs were considered, and we also removed genes labeled as “dubious”, mitochondrial or 2-micron plasmid encoded genes, resulting in a set of 5,894 genes.

For homology detection we used fourteen genomes from the *Saccharomycetales* order, see Table 1. Five of these genomes belong to the *Saccharomyces* genus. All genomes are fully sequenced with an assembly status equal to “contigs”, “scaffold”, “chromosomes” or “complete genome”. Full genome sequences were obtained from NCBI Genome Project, while proteomes were downloaded from Uniprot [35]. It should be noted that many better annotated fungal genomes are available today. However, we choose to use this set, as it is representative of what was used in earlier studies.

**Table 1.**
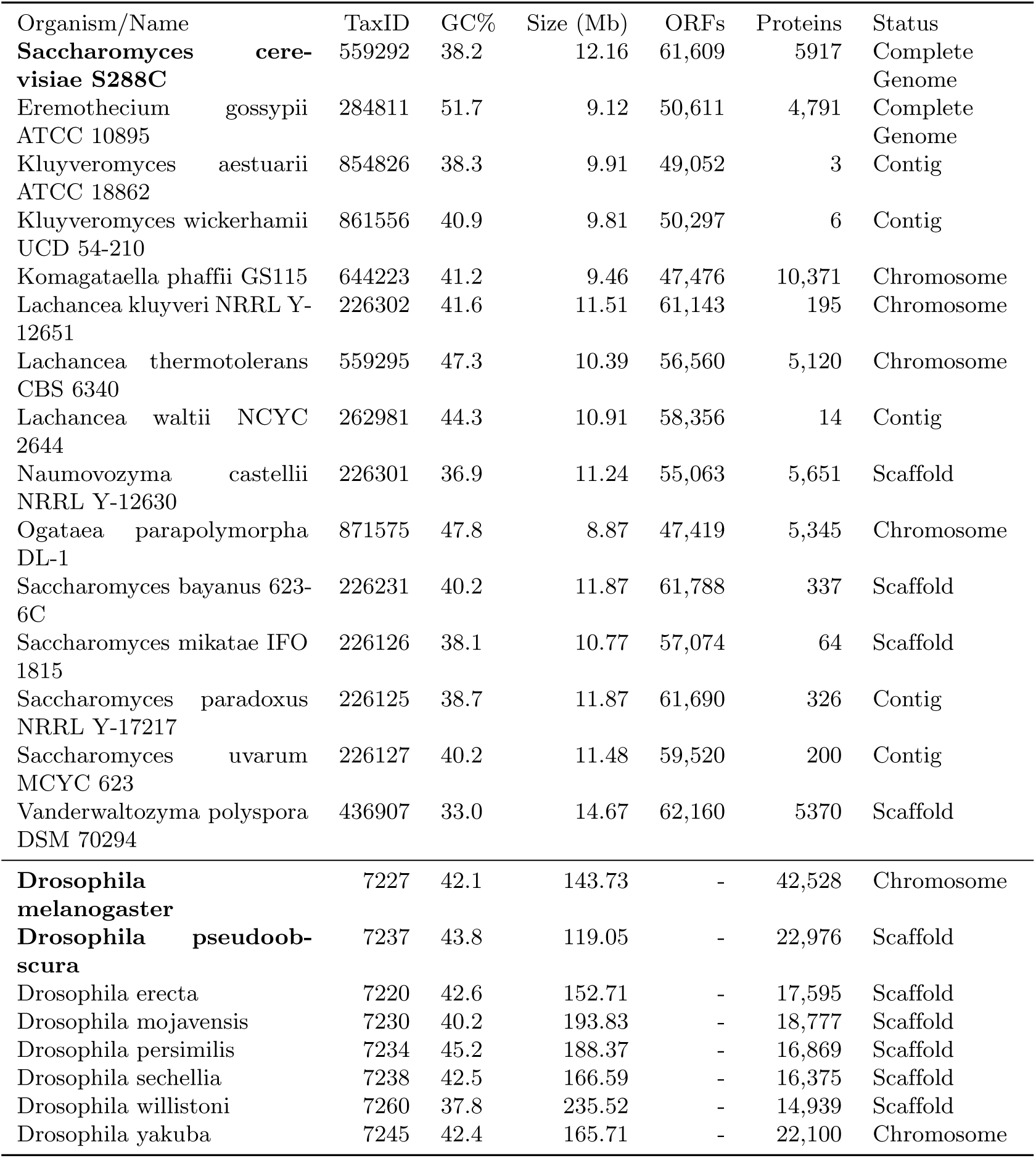
Summary of the genomes used in this study divided into 15 Ascomycetes and 8 Drosophila genome. GC% = Guanosyne+Cytosine content of the entire genome; ORFs = number of Open Reading Frames; Proteins = number of annotated proteins in UniprotKB. Estimates of number of chromosomes were obtained from Keogh et al. [54] when not present in NCBI data.

#### Estimation of evolutionary distances between genomes

To estimate the phylogenetic distance between the genomes we used the amino acid sequence of the well-conserved gene mre11 (nuclease subunit of the MRX complex, involved in DNA repair) as a proxy for evolutionary distance. A multiple sequence alignment was created using Clustal Omega [36] with default parameters. Thereafter, a phylogenetic tree was built using the Maximum Likelihood (ML) algorithm implemented in PhyML [37], version 20160207. The estimated number of substitutions per site, using the LG model [38] then represents the evolutionary distance.

#### RNA-seq data

RNA-Seq expression data for *Saccharomyces cerevisiae* was obtained from two studies [39, 40]. The first study is from the wild-type sample, while the second dataset consists of three replicates of samples with chromosome X disomy and three replicates of the wild type haploid. We used expression in any of the studies as evidence for gene expression.

### Drosophila data

Drosophila genomes were downloaded from modENCODE [41], see Table 1. In addition 33 RNA-Seq datasets for Drosophila melanogaster and 32 datasets for Drosophila pseudoobscura were used to study the expression. The RNA-Seq experiment includes expression from different stages of the life cycle. All the datasets were processed using STAR [42]. FeatureCounts was used to obtain the number of successfully assigned reads [43].

### Homology identification

To detect orphans it is necessary to use a search tool to infer homology between genes. The search can be performed using a database of proteins, or complete genomes. Searching a full genome can be done using either the nucleotide sequence, or by using a six-frame translation of all ORFs in a genome. Further, different search methods can be used, including BLAST [29] and HMMER [44].

In this study we used three variations of BLAST [29]. First, nucleotide-blast (blastn) was used to detect homology between a gene in the query genome and another genome. Secondly protein-blast (blastp) was used to search for homology between annotated proteins. Finally, tblastn, that searches a protein sequence against 6-frame translated nucleotide database, was used. For all BLAST variants NCBI BLAST+ version 2.6.0 was used with default parameters.

To detect potentially missed homologies with other genomes we searched the final set of orphans against the NCBI NonRedundant (nr) database. Here, we used three BLAST variants for *S. cerevisiae* (blastp, blastn and tblastn), and two for *Drosophila* (blastp and tblastn).

To estimate the number of false positives, i.e. genes incorrectly labelled as orphans, we scrambled the sequences of the 5894 yeast proteins. These were searched against the database using the same methods and parameters as the regular queries. Results are reported for different E-value cutoffs in Table S9.

### Comparison of orphan sets

To analyze the similarities between sets of orphans, we used the *F*_1_ score, that normally is used to classify correct and incorrect predictions:

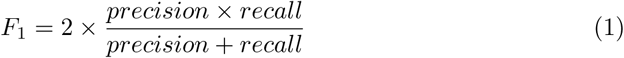

Where

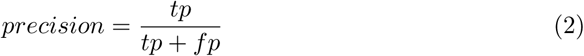

And

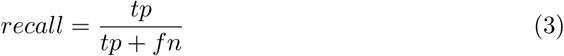

with *tp* = true positives, *fp* = false positives and *fn* = false negatives. Here, we do refer genes present in both sets as true positives, while false positives are those present in the first but not in the second and false negatives are those present in the second but not in the first. Thus, a F1 score of 1 denotes two identical sets.

### Identified orphans

In supplementary tables S1, S2 and S3 the list and sequences of the identified orphans are shown for all three species.

## Results and Discussion

Identification of orphans is dependent on the method used to detect homology and what database is searched. We used 5,894 *Saccharomyces cerevisiae* protein coding genes and searched a database consisting of fourteen *Saccharomycetales* genomes at different evolutionary distance from *Saccharomyces cerevisiae*, see Table 1.

These yeast genomes are similar in size (9-12 Mb), but the number of annotated proteins varies significantly. Obviously, the number of proteins does not correspond to the real size of the proteomes; rather, it reflects the annotation status of these genomes. In particular, it is apparent how *Saccharomyces paradoxus*, a close relative to *S. cerevisiae*, is poorly annotated despite being fully sequenced. Besides representing the status of earlier studies, this also constitutes a good example to illustrate how different levels of annotations influence the results. This choice of genomes might also be representative for other less studied lineages, where well-annotated genomes is lacking.

### Homology detection strategies

Three different search strategies were used to detect homologs of 5894 *S cerevisiae* genes. To exemplify the effectiveness of different homology detection strategies we use an E-value ≤ 0.001 and report the average number of genomes were a hit was found. This means that each gene can have a maximum of 14 hits. Obviously it would be assumed that the vast majority of genes should have at least one homolog in all fourteen genomes. (i) With blastn on average 2.6 hits per gene were found, indicating that many homologous relationships are missed. (ii) Using blastp increased the average number of hits to 5.8. The main reason why no hits are found in about half the genomes is due to the poor status of gene annotations, as discussed above. (iii) Using tblastn, i.e. a six-frame translation of the genomes, 12.8 hits per gene were found. This means that on average the genes in *Saccharomyces cerevisiae* have hits in all but one of the target genomes. Clearly, only the last strategy finds most homology relationships.

### Number of orphans

In Table 2 we report the number of the genes classified as orphans using the different homology search methods and at different E-value cutoffs. It can be seen than the search methodology has a much bigger effect that the cutoff used. Therefore, the fact that previous studies used different E-values, from 0.1 in Lu et al. [28] to 0.001 in Ekman et al [4], cannot explain the differences in their results. Using blastn or blastp, hundreds of proteins are classified as orphans independently of the cutoff used, while using tblastn the number of orphans is much lower. This shows that using blastp or blastn results in an overestimation of the number of orphans.

**Table 2.** Orphans detected in *S. cerevisiae* by different methods and by their intersection (“Shared”) at different E-value cutoffs. The E-value chosen for this study is highlighted in boldface.

| e-value | blastn | blastp | tblastn | tblastn+blastp | Shared |
| --- | --- | --- | --- | --- | --- |
| 0.10000 | 701 | 261 | 52 | 45 | 44 |
| 0.01000 | 701 | 280 | 63 | 56 | 55 |
| **0.00100** | **701** | **291** | **72** | **63** | **61** |
| 0.00010 | 701 | 304 | 75 | 66 | 62 |
| 0.00001 | 701 | 312 | 81 | 74 | 68 |

For tblastn the number of orphans is approximately halved when an E-value cutoff of 0.1 is used compared with 10^−5^. For simplicity, we decided to use an E-value cutoff (0.001) similar to earlier studies. At this E-value threshold, the number of false positive orphans (i.e. the number of scrambled sequences deemed as orphans, see Methods) is 2 with blastp and 1 with tblastn (Table S9). Using tblastn with this cutoff, 72 orphans are identified. However, the tblastn hits do not constitute a strict subset of the ones found by blastp. Therefore, using a combined strategy of blastp+tblastn reduces the number of orphans to 63 orphans are found by all methods. We will refer to this as our “strict orphan set”.

### Comparison with previous estimates of orphans in *S. cerevisiae*

We compare the different sets of orphans identified in *S. cerevisiae* with earlier studies. In Ekman et al. [4] a set of 157 *S. cerevisiae* specific proteins was identified by using a method based on blastp: each amino acid in the query proteins was assigned to one of six phylogenetic levels. The lowest level corresponds to those found exclusively in *S. cerevisiae*. If all the amino acids of a protein are assigned to this level, the protein is classified as an orphan. Of the 188 proteins found in this way, 31 were removed in a second step, after searching for more distant homologs with HMMER [44] in the Pfam database [45], and with HHsearch [46] in other databases. Two proteins from the Ekman set are not present in our study, resulting in an overlap of 155 proteins.

In Carvunis et al. [5], the complete set of *S. cerevisiae* ORFs was divided into groups of progressively higher conservation, based on blast hits in a set of 14 other yeast genomes. Three blast variants were used (blastp, tblastx, tblastn). The set of 143 genes called ORFs_1_ (i.e. the least conserved) should be analogous to *S. cerevisiae* species-specific orphans as we define them here. However, in this study ORFs labeled as “dubious” were also included. Thus, only 34 of the 143 genes are present in our dataset, see above. 30 out of the 34 orphans from Carvunis are also present among the orphans from Ekman, showing the more conservative estimates by Carvunis.

In Lu et al. [28], a set of 22,793 *S. cerevisiae* CDS is considered. By studying their conservation within the *Saccharomycetaceae* clade, and, crucially, detecting the conserved synteny by using tblastn, they propose a set of 4340 *S. cerevisiae*-specific genes, of which 47 are annotated as orphans. However, only 16 of these are present in our dataset; only 6 of these 16 genes are present in the 61 strict orphans and none of these are present in the Carvunis set.

To compare all these orphans sets we calculated the *F*_1_ score (see Methods) for each pair of methods, see Table 3 and Figure 1. The Ekman set is most similar to blastp, with a *F*_1_ score of 0.65, while the Carvunis set is close to the tblastn set (*F*_1_ ∼ 0.57). This is in agreement with the methodologies used, as Ekman searched proteomes using blastp, and Carvunis full genomes using tblastn.

**Table 3.** Agreement (expressed as *F*_1_ score, see Methods) between the results with our methods and the *S. cerevisiae* orphan sets proposed by Ekman et al., Carvunis et al. and Lu et al. For each pair of orphan sets, higher F1 score represents a larger fraction of genes present in both sets.

| Method | Ekman | Carvunis | Lu |
| --- | --- | --- | --- |
| tblastn+blastp | 0.55 | 0.62 | 0.15 |
| tblastn | 0.56 | 0.57 | 0.14 |
| blastp | 0.65 | 0.20 | 0.10 |
| blastn | 0.24 | 0.08 | 0.03 |
| Ekman | 1.00 | 0.32 | 0.14 |
| Carvunis | 0.32 | 1.00 | 0.00 |
| Lu | 0.14 | 0.00 | 1.00 |

**Fig 1.**
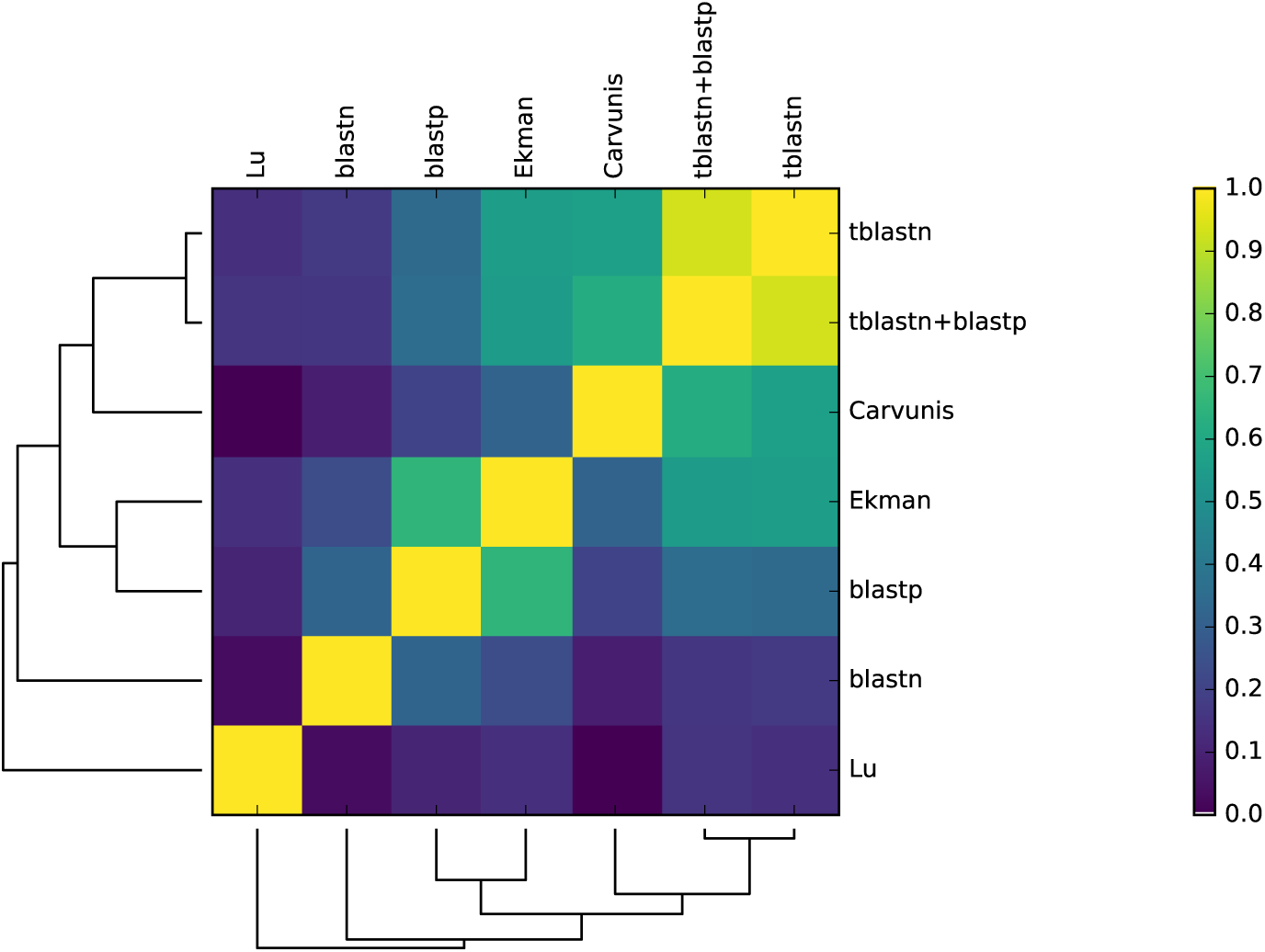
Agreement (expressed as *F*_1_ score) of sets of *S. cerevisiae* orphans obtained with the different methods and proposed by Ekman et al., Carvunis et al. and Lu et al.

Out of the 61 strict orphan genes, 58 are also present in the Ekman orphan set, and 30 in the Carvunis set, see Figure 2. However, only 6 of these orphans are present in the Lu set, which is a clear outlier.

**Fig 2.**
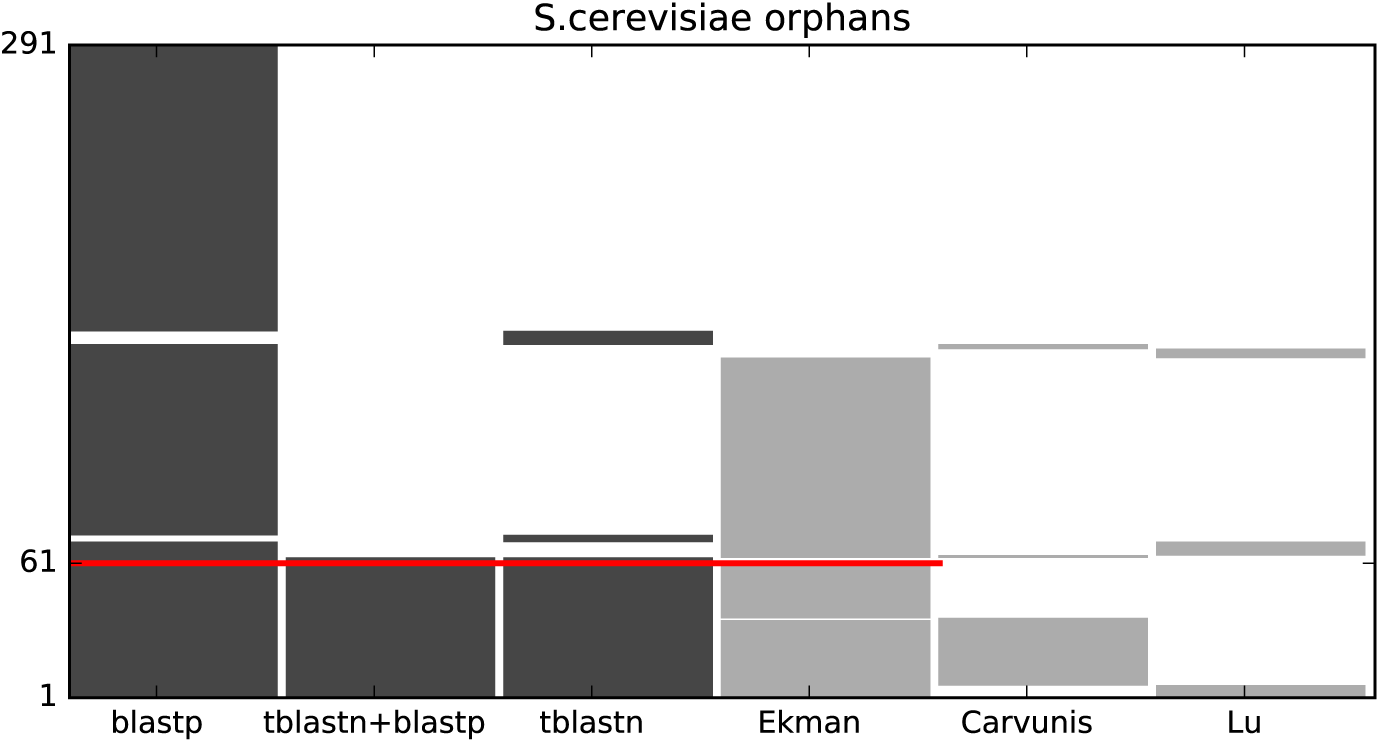
Genes defined as orphans in *S. cerevisiae* by the different methods, in the subset of genes deemed as orphans by at least one method. A red line marks the 61 genes defined as orphans by all methods (strict orphans). Sets of orphans proposed by previous studies (Ekman et al, Carvunis et al., Lu et al.) are shown in light grey.

### Effect of number of species and evolutionary distance

The number of proteins classified as orphans is strongly dependent on the genomes included in the search for homologs. To evaluate this effect we explored how altering the number of species affects the number of identified orphans. We estimated the orphans by using one single species at the time, and using blastp+tblastn as a homology detection method.

Clearly, there exists a strong relationship between the evolutionary distance of the species and the number of proteins classified as orphans, see Figure 3. Whenever *S. paradoxus* is not included the number of genes classified as orphans is more than 156, and when all *saccharomyces sensu stricto* genomes are ignored the number is over 397.

**Fig 3.**
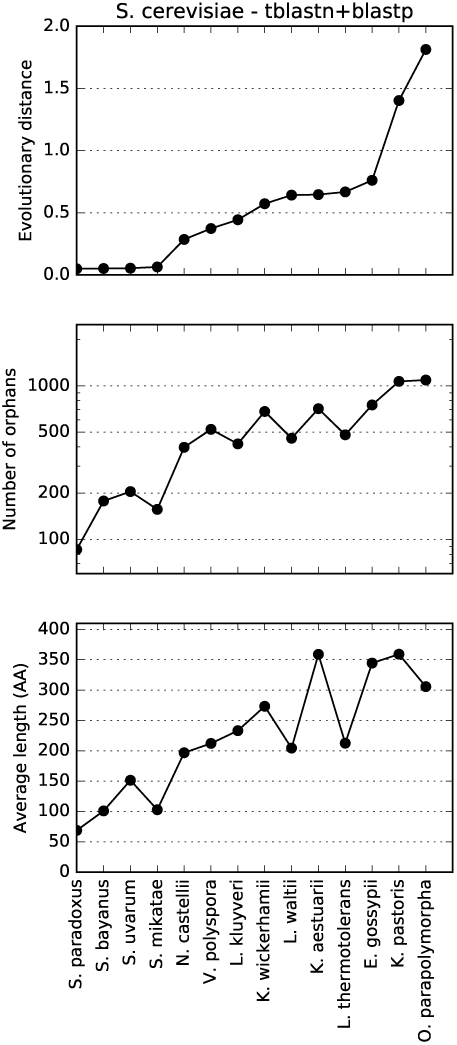
Evolutionary distance, number and average length of orphans in *Saccharomyces cerevisiae* using tblastn+blastp, estimated with one neighbor species at a time.

It can also be noted that including a single closely related species is not sufficient as the loss of genes is frequent. The number of orphans classified when only using *S. paradoxus* is over 85 compared to less than 67 when five or more of the closest species are included (data not shown).

### Properties of the different orphan sets

Several studies have used sets of orphans to draw conclusions on the mechanisms of *de novo* creation of proteins [27, 47–49]. However, the number of proteins identified as orphans depends on the homology detection method and the database used. The most stringent methods identify 34-52 orphans in *S. cerevisiae*, while other methods identify many more. Knowing which of these genes really are *de novo* created or even functional is problematic [33]. Therefore, it is difficult to know which method is the most accurate. However, to gain some insights we study two properties of the different sets, the average protein length and the fraction of intrinsically disordered residues using IUPred [50]. It is in generally agreed that orphan genes should be short, while intrinsic disorder among orphan proteins might be a signal of fast evolution.

It has been previously noted that orphan genes tend to be short. This has been attributed to the *de novo* gene creation mechanism, in which a short ORF gets translated, and becomes longer as it gets fixed in the population [5]. Among our different orphan sets, the orphans are on average shorter than other genes, and there is actually a strong correlation (Cc=0.90) between number of orphans and length, see Table 4.

**Table 4.** Number of orphans found in *S. cerevisiae* by different methods, their average length in amino acids and their intrinsic disorder (predicted using IUPred long).

| Method | Number | Length(AA) | Disorder(%AA) |
| --- | --- | --- | --- |
| tblastn+blastp | 63 | 54 | 3.8% |
| tblastn | 72 | 55 | 8.7% |
| blastp | 291 | 127 | 10.3% |
| blastn | 701 | 254 | 15.8% |
| Carvunis | 34 | 62 | 3.1% |
| Ekman | 155 | 72 | 5.7% |
| Lu | 16 | 77 | 5.6% |

Intrinsic disorder has also been associated with orphan genes [27]. We predicted the intrinsic disorder using IUPred [50] and also here there is a strong correlation (Cc=0.91) between number of orphans and the fraction of residues predicted to be disordered. In the orphan set from the most restrictive method, the fraction of disordered amino acids is 3.8%, while it is 16% using the least stringent methods. This indicates that if a loose criterion is used to identify orphans they will appear to be more disordered.

It is well established that disordered proteins have larger variation in length [26] and are fast evolving [51]. Taking this into account, a picture emerges where less sensitive methods could miss homologies to fast-evolving disordered proteins. This causes orphans detected by such methods to appear longer and more disordered than *de novo* created genes. By using the more sensitive strategy proposed here, these incorrect assumptions could be avoided.

### Identification of potential false positives

The 63 orphans identified by blastp+tblastn were studied more carefully. To ensure that no homologs exist to these genes, we searched in several other databases using different methods: PSI-BLAST against the NCBI Environmental Protein Sequences Database (env nr), hhblits against Uniclust30 (April 2017) and HMMER-hmmscan against PFAM. None of these searches produced any significant hit outside *S. cerevisiae*.

However, when searching the GenBank NonReduntant (nr) database some hits are found using blastn or tblastn, see Table 5. Using tblastn four significant hits appear; two of these are to a bacteria (Bacillus), one to *Saccharomyces paradoxus* and one to *Saccharomyces boulardii*, a fungal genome not present in our set of genomes. The bacterial hits are either rare examples of lateral transfer or due to incorrect entries in the database, possibly obtained by some contamination.

**Table 5.**
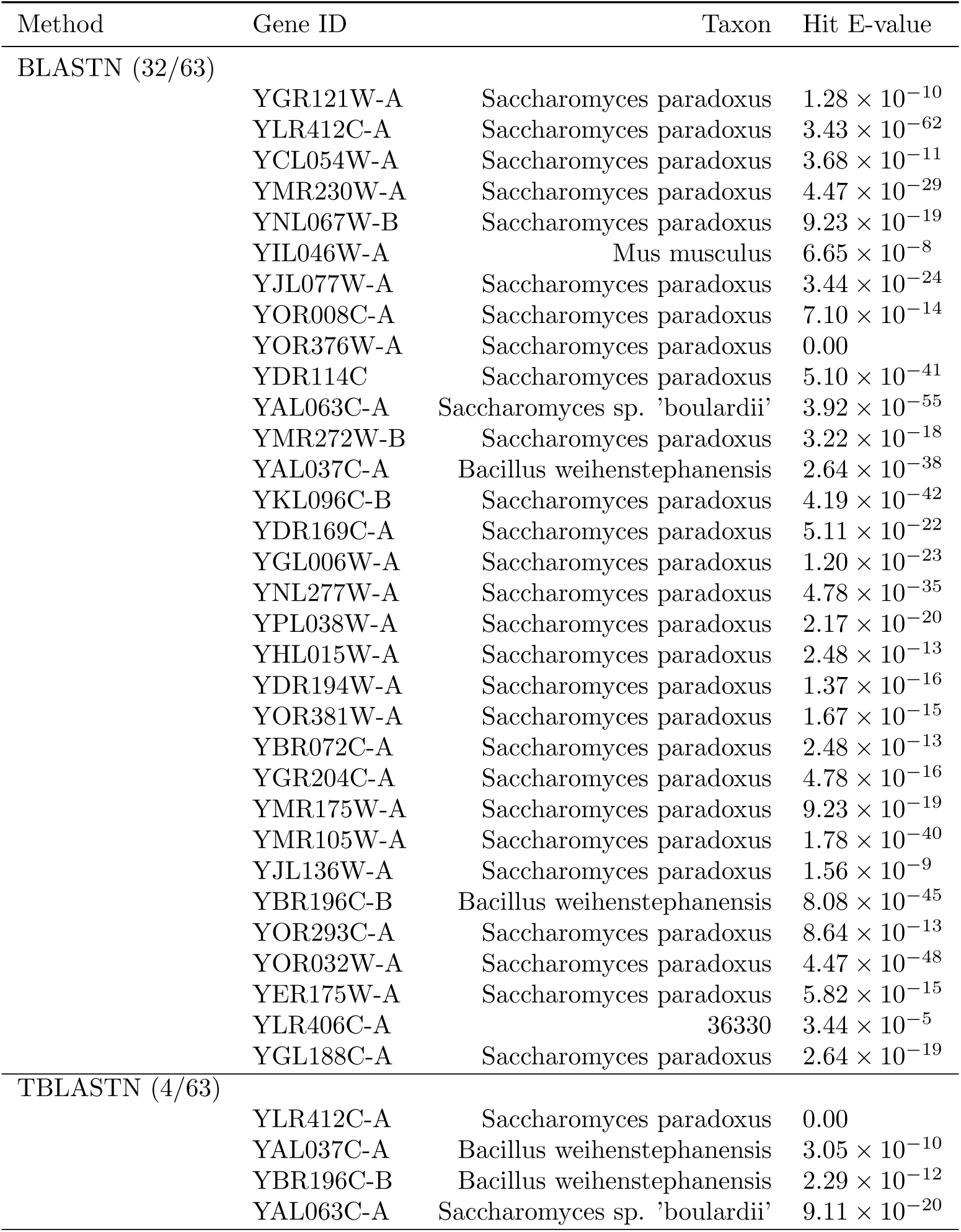
63 genes deemed as orphans in *S. cerevisiae* by blastp+tblastn, are searched in GenBank NonRedundant (nr) database, using three BLAST variants: blastp, blastn and tblastn. No hits are found with blastp. For each of these searches, it is reported the number (in parentheses) of queries with at least one hit with E-value < 0.001. For each gene, the E-value of its best hit in nr and the taxon of the target species are shown.

| Method | Gene ID | Taxon | Hit E-value |
| --- | --- | --- | --- |
| BLASTN (32/63) |  |  |  |
| | YGR121W-A | Saccharomyces paradoxus | $1.28 \times 10^{-10}$ |
| | YLR412C-A | Saccharomyces paradoxus | $3.43 \times 10^{-62}$ |
| | YCL054W-A | Saccharomyces paradoxus | $3.68 \times 10^{-11}$ |
| | YMR230W-A | Saccharomyces paradoxus | $4.47 \times 10^{-29}$ |
| | YNL067W-B | Saccharomyces paradoxus | $9.23 \times 10^{-19}$ |
| | YIL046W-A | Mus musculus | $6.65 \times 10^{-8}$ |
| | YJL077W-A | Saccharomyces paradoxus | $3.44 \times 10^{-24}$ |
| | YOR008C-A | Saccharomyces paradoxus | $7.10 \times 10^{-14}$ |
|  | YOR376W-A | Saccharomyces paradoxus | 0.00 |
| | YDR114C | Saccharomyces paradoxus | $5.10 \times 10^{-41}$ |
| | YAL063C-A | Saccharomyces sp. 'boulardii' | $3.92 \times 10^{-55}$ |
| | YMR272W-B | Saccharomyces paradoxus | $3.22 \times 10^{-18}$ |
| | YAL037C-A | Bacillus weihenstephanensis | $2.64 \times 10^{-38}$ |
| | YKL096C-B | Saccharomyces paradoxus | $4.19 \times 10^{-42}$ |
| | YDR169C-A | Saccharomyces paradoxus | $5.11 \times 10^{-22}$ |
| | YGL006W-A | Saccharomyces paradoxus | $1.20 \times 10^{-23}$ |
| | YNL277W-A | Saccharomyces paradoxus | $4.78 \times 10^{-35}$ |
| | YPL038W-A | Saccharomyces paradoxus | $2.17 \times 10^{-20}$ |
| | YHL015W-A | Saccharomyces paradoxus | $2.48 \times 10^{-13}$ |
| | YDR194W-A | Saccharomyces paradoxus | $1.37 \times 10^{-16}$ |
| | YOR381W-A | Saccharomyces paradoxus | $1.67 \times 10^{-15}$ |
| | YBR072C-A | Saccharomyces paradoxus | $2.48 \times 10^{-13}$ |
| | YGR204C-A | Saccharomyces paradoxus | $4.78 \times 10^{-16}$ |
| | YMR175W-A | Saccharomyces paradoxus | $9.23 \times 10^{-19}$ |
| | YMR105W-A | Saccharomyces paradoxus | $1.78 \times 10^{-40}$ |
| | YJL136W-A | Saccharomyces paradoxus | $1.56 \times 10^{-9}$ |
| | YBR196C-B | Bacillus weihenstephanensis | $8.08 \times 10^{-45}$ |
| | YOR293C-A | Saccharomyces paradoxus | $8.64 \times 10^{-13}$ |
| | YOR032W-A | Saccharomyces paradoxus | $4.47 \times 10^{-48}$ |
| | YER175W-A | Saccharomyces paradoxus | $5.82 \times 10^{-15}$ |
| | YLR406C-A | 36330 | $3.44 \times 10^{-5}$ |
| | YGL188C-A | Saccharomyces paradoxus | $2.64 \times 10^{-19}$ |
| TBLASTN (4/63) |  |  |  |
|  | YLR412C-A | Saccharomyces paradoxus | 0.00 |
| | YAL037C-A | Bacillus weihenstephanensis | $3.05 \times 10^{-10}$ |
| | YBR196C-B | Bacillus weihenstephanensis | $2.29 \times 10^{-12}$ |
| | YAL063C-A | Saccharomyces sp. 'boulardii' | $9.11 \times 10^{-20}$ |

When using blastn, hits are found for about half of the orphans, while using blastp no hits are found, see Table 5. Almost all hits are with *Saccharomyces paradoxus*. Upon closer inspection these hits are with regions of the genome not annotated to contain Open Reading Frames. Earlier it has been shown that young ORFs in *S. cerevisiae* correspond to regions in the closely related *S. paradoxus* that either lack a start codon, or contain a premature stop codon [28]. Therefore, it is possible that either these regions in *S. paradoxus* correspond to regions for *de novo* gene creation in *S. cerevisiae* or that they became pseudo-genes in *S. paradoxus*. However, given the poor status of annotations in *S. paradoxus* makes it also quite likely that these hits actually represent functional genes in *S. paradoxus* as well. To distinguish these scenarios would require a more detailed analysis.

### Transcriptional evidence

We have investigated the presence of the orphans we identified in several datasets of RNA-Seq experiments. These results are shown in Table 6 and Table S4. 86% of the classified orphans show transcriptional evidence, compared to the 98% of all other genes. This indicates that most of the identified orphans, despite being young and *de novo* created, have an active role in the cell.

**Table 6.** For *S. cerevisiae, D. melanogaster* and *D. pseudoobscura* it is shown the fraction of orphan and ancient proteins that are found in RNA-Seq experiments, at different E-value cutoffs.

| E-value | *S. cerevisiae* |  | *D. melanogaster* |  | *D. pseudoobscura* |  |
| --- | --- | --- | --- | --- | --- | --- |
|  | Orphans | Ancient | Orphans | Ancient | Orphans | Ancient |
| 0.10000 | 0.82 | 0.98 | 0.46 | 0.51 | 0.38 | 0.53 |
| 0.01000 | 0.86 | 0.98 | 0.46 | 0.51 | 0.43 | 0.52 |
| 0.00100 | 0.86 | 0.98 | 0.46 | 0.51 | 0.42 | 0.52 |
| 0.00010 | 0.86 | 0.98 | 0.49 | 0.51 | 0.42 | 0.52 |
| 0.00001 | 0.88 | 0.98 | 0.49 | 0.51 | 0.43 | 0.52 |

### Orphans in Drosophila

We also examined orphan classification in *Drosophila*, another well-studied clade. We identified orphans by using tblastn, blastp and a combination of them for *D. melanogaster* and *D. pseudoobscura*, see Table 7.

**Table 7.**
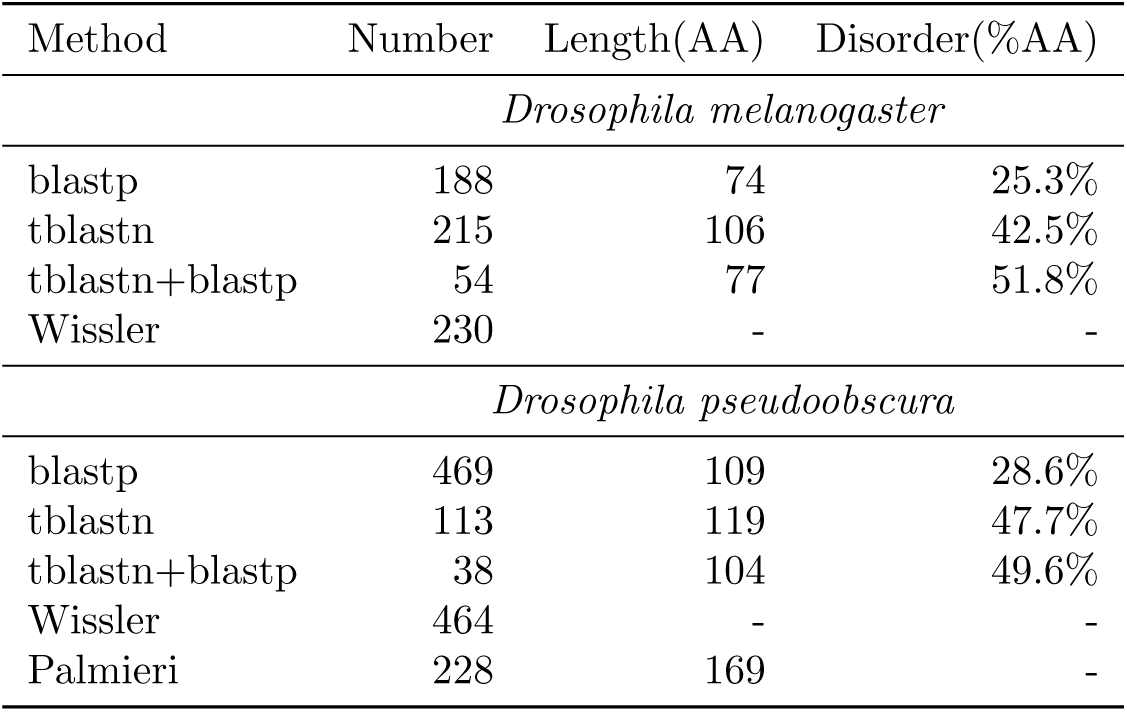
Number of orphans found by different methods in *Drosophila melanogaster* and *Drosophila pseudoobscura*, their average length in amino acids and their intrinsic disorder (predicted using IUPred long).

| Method | Number | Length(AA) | Disorder(%AA) |
| --- | --- | --- | --- |
| *Drosophila melanogaster* |  |  |  |
| blastp | 188 | 74 | 25.3% |
| tblastn | 215 | 106 | 42.5% |
| tblastn+blastp | 54 | 77 | 51.8% |
| Wissler | 230 | - | - |
| *Drosophila pseudoobscura* |  |  |  |
| blastp | 469 | 109 | 28.6% |
| tblastn | 113 | 119 | 47.7% |
| tblastn+blastp | 38 | 104 | 49.6% |
| Wissler | 464 | - | - |
| Palmieri | 228 | 169 | - |

Wissler et al. [52] proposed 230 orphans in *D. melanogaster* and 464 in *D. pseudoobscura*, while Palmieri et al. [53] estimated 228 orphans in *D. pseudoobscura*. These estimates are similar to blastp alone (188 in *D. melanogaster* and 469 in *D. pseudoobscura*). Unfortunately, a direct comparison of the sets is difficult as the gene identifiers from the two earlier studies are not compatible with Uniprot.

The combined method identifies only 54 and 38 orphans in *D. melanogaster* and *D. pseudoobscura* respectively. In both species the number of orphans identified by the combined method is clearly lower then for the individual methods. This difference is larger than in *S. cerevisiae*. This can be explained by difference in genome size and the less gene-dense drosophila genomes. It is clear that the blastp sets of orphans do include many proteins with homologs detectable by tblastn, just as the Ekman set for *S. cerevisiae*.

It can be noticed that 17 out of 54 (31%) of *D. melanogaster* orphans have hits in nr when using blastp, mainly in *Drosophila simulans*; but only 2 out of 38 (5%) of *D. pseudoobscura* orphan proteins have hits in nr, see Tables S7 and S8. In *Drosophila melanogaster* we found transcriptional evidence for roughly half of all the genes and for 46% of the orphans see Tables S5. In *Drosophila pseudoobscura*, 50% of all the genes in our dataset are found to have evidence for expression and in its orphans the fraction rises to 42% see Tables S6. The lack of transcriptional evidence for most of orphan and non-orphan genes in these two organisms makes it difficult to evaluate if the orphans are functional or not.

## Conclusions

Here, we show that performing homology searches on a handful of closely related, six-frame translated genomes is vital to determine a correct set of orphan genes. If this is done accurately, the set of orphans should represent *de novo* created genes. However, if this is not done properly incorrect conclusions on their properties might be drawn. In particular, orphans might appear longer and more disordered than they really are. We propose that a combination of blastp and tblastn is more powerful than any of these methods alone. This advantage is particularly evident in higher eukaryotes such as *Drosophila*, where the number of orphans drops from hundreds to 54 in *D. melanogaster* and 38 in *D. pseudoobscura*. However, it should be kept in mind that the annotation status of many genomes is still partial; hits provided by tblastn might also correspond to non-functional ORFs. We believe that to this day the estimate of orphans in higher eukaryotes might have been affected by the gene annotations and the choice not to search complete genomes using their six-frame translations.

## Acknowledgments

This work has been supported by the Swedish Research Council (VR-NT 2012-5046 and 2016-06301) and the Swedish E-science Research Center.

## Supporting Information Legends

**Table S1.** List and sequence of orphans found in *S. cerevisiae* in fasta format.

**Table S2.** List and sequence of orphans found in *D. Melanogaster* in fasta format.

**Table S3.** List and sequence of orphans found in *D. pseudoobscura* in fasta format.

**Table S4.** For the 52 *S. cerevisiae* genes out of the 63 orphans found by blastp+tblastn, it is shown the expression level in 7 different RNA-Seq experiments. The values in the columns corresponds to the RNA expression data from the two experiments taken into consideration. The values in the first column (RNA seq exp 1) are the log2 tag count reported in [39]. The values in the columns RNA seq exp 2, RNA seq exp 3 and RNA seq exp 4 are the Reads Per Kilobase Million (RPKM) values for the three replicates of the Disome 10 (chromosome X disomy) reported in [40]. The values in the columns RNA seq exp 5, RNA seq exp 6 and RNA seq exp 7 are the RPKM values for the three replicates of the wild type haploid reported in [40].

**Table S5.** For the 25 *D. melanogaster* genes out of the 54 orphans found by blastp+tblastn, it is shown the expression level in 33 different RNA-Seq datasets. The values in the columns corresponds to the RNA expression data in Reads Per Kilobase Million (RPKM).

**Table S6.** For the 16 *D. pseudoobscura* genes out of the 38 orphans found by blastp+tblastn, it is shown the expression level in 32 different RNA-Seq datasets. The values in the columns corresponds to the RNA expression data in Reads Per Kilobase Million (RPKM).

**Table S7.** 54 genes deemed as orphans in *Drosophila melanogaster* by blastp+tblastn, are searched in GenBank NonRedundant (nr) database, using two BLAST variants: blastp and tblastn. For each of these searches, it is reported the number (in parentheses) of queries with at least one hit with E-value < 0.001. For each gene, it is shown the E-value of its best hit in nr, and the target species.

**Table S8.** 38 genes deemed as orphans in *Drosophila pseudoobscura* by blastp+tblastn, are searched in GenBank NonRedundant (nr) database, using two BLAST variants: blastp and tblastn. For each of these searches, it is reported the number (in parentheses) of queries with at least one hit with E-value < 0.001. For each gene, it is shown the E-value of its best hit in nr, and the target species.

**Table S9.** False positive orphans found but blastp and tblastn at different E-values thresholds. These are the hits obtained using as queries scrambled sequences of the original ones.

## Notes

#### Summary of Updates

Updated version for revision 2

