## Supplementary material for "The classification of orphans is improved by combining searches in both proteomes and genomes"

Walter Basile[1,2], Marco Salvatore[1,2], Arne Elofsson[1,2,3,*],

**1 Science for Life Laboratory, Stockholm University SE-171 21 Solna, Sweden**
**2 Department of Biochemistry and Biophysics, Stockholm University, SE-106 91 Stockholm, Sweden**
**3 Swedish e-Science Research Center (SeRC)**

### Supporting Information

```
>YGR121W-A
MSRHKYKLWMCIAKGKRGVGERNTFVPKCSYAPFCASEVLDQLLRGDNSIVYNFYLKNGSGSVYIGSVFLI*
>YLR157W-E
MIVDFYSNTLRHCETLRSQPCSLFSSLYARSFQSSCTLHVAEPSPGFHMYGCHT*
>YOL038C-A
MKYMGSFLRKAATTNLFNSIKKRKVQNRAMS*
>YLR157W-D
MKFQYALAKEQLGSNSRSGVKKLISKHHWLPEYYFSDLSFSVVQQWDSRAIEKTTIISCMRPANQEIYPL*
>YLR412C-A
MHLCQNGHYYKPHRASAEKVPYLKKKKKNSRNEGKAKKKNEKKKIGTVEFFQQKKKEKKRVLNAVCGL*
>YCL054W-A
MIRQKIFVFIVKSRRNSICPAIRRKEDY*
>YJL136W-A
MTRCISKKMLLEVDALSLIYSPHLYMS*
>YMR230W-A
MCIIVKYSQCNLFFQDVLRCRHVLCMLFSIVVHFRLSSFVSVCGVRIMCNCINCSRFIHNIN*
>YNL067W-B
MCKLMWCTGVVSKTALLTGNFFFSSSEFFFKATHRKSENYLNGRQT*
>YIL046W-A
MMCVCIPKKKLMDWRVYYIYSYVVCLYMCGSDCACICVLACVVQCVCFNVEMRL*
>YHR214C-D
MEDHTLVAIVVFFGNGEPFHVSLSVEMVFVLLLSSTRIHEVVVLICYKLQHATWSWGNMSKNFSLKPDISLSFLLDIISINDICIYGCIALTVVFIL*
>YGR174W-A
MNLNAYFEAYQAIFPFLLEAFLRKEQKV*
>YAL064W
MNPFASLEGQDNISSVFFLHMQQFESQVKDRFRFPIFRLERKTFGNSCYQVETLKVKCRPRHAKSCNLLTLLFKSRTQSVLVPNFGFLILNSEP*
>YJL077W-A
MPGIAFKGKDMVKAIQFLEIVVPCHCTT*
>YOR008C-A
MWRSYLVFLFFMTPRIQTYCPVPVLRSMAVLNIISPLIIFVSPIKKQDSLHSSACYANLTLVEKLQLWHSMSND*
>YOR376W-A
MTFHLGWTILWYNQAYLEVWATVFQDEMHKYSLHPQSRDAKTKFCCCIPFK*
>YDR114C
MKTFFLIEAGRALQIVAFRPAITTVLFQRFSVSFSCSYFTCCTSQLLRENWLFKFLTAFDHVIRASNDLSTMRFMIMYIYVYIYIYTVLRKRLSCYMLIL*
>YLR156W
MKFQYALAKEQLGSNSRSGVKKLISKHHWLPEYYFSDLSFSVVQQWDSRAIEKTTIISCMRPANQEIYPLRHCETLRSQPCSLFSSLYARSFQSSCTLHVAEPSPGFHMYGCHT*
>YAR035C-A
MRLNYSRCYYSSQRRRQSLPKRFPLI*
>YAL063C-A
MCPRTVLLIININHWFYDKNIVRIILTFRLDSGHISDICFINKNLANALITADISLLKRHDIRCTKYIITYYQRYRNKEKGKFISLCKNTIISSSV*
>YMR272W-B
MRSLVFVQLSLLSWEIFCGERSFVSMKAIFSCMYV*
>YBR296C-A
MKVLDDWFSRKFSKAVHGNNHGTISLSTLSYIRVHKLVK*
>YAL037C-A
MSISFPKMQHLIVMTTIGDKKVNNNIILFL*
>YOR293C-A
MKLLFLNIIVVRRHLHCKSYRLSPWYIYIYGDYLLYTTEIPYKPFTRQP*
>YGL007C-A
MLPSISFDYIKRPNIVLFSNVLSLSSNI*
>YMR247W-A
MAHKCASAKLLSGIMALLFNGKSLLRPICLHVHNHLVSNSDTNIVWP*
>YBR182C-A
MKIFTLYTMIQQYFFDNGGVYSIKNFYSAVPKEKMNIILVSLDCELQKLALKLSKKTSGTHTTH*
>YIR018C-A
MPSDYTSHYPVILIKKKKKKIAGMYRHSKRYLEIMSTASAQFVGN*
>YGL006W-A
MLIFIIHYHRHLALHLMGAFQKHSNSISPPPRKGFI*
>YNL277W-A
MTLAYYGQPVKMCHILPPLRSLPVLVGKKKLKKKKSQTTNNHVIFLFTLFIKLLKTHNRMSL*
>YOL155W-A
MFYGSFNKCVTGYSCRMAIHYYVYRIIKSATRPDYKSNTQILVL*
>YPL038W-A
MVCRFVHHSRVIFISIYDFLSTKGKKNMYNYTQEKKTKQKNTFTQASIYYENFFESYRTISCL*
>YBR200W-A
MLLCFHMCQRIMWLPFDLMKWRRFHCGAVLVGTLSLRNRSPKILSLYFISDRTG*
>YDR194W-A
MKQMMIEASISKDTLRLLICFFEIKQCHISLQPTCYYQNWVRYSSIYYQL*
>YHR022C-A
MKIKFSRGARFSATFSFDKYPFLLYEVVR*
>YOR381W-A
MFNVALPSQDKKSEFYKQKRLSLRTNILDEFLRVAKTQDNRQNNFVSQYRNDEIY*
>YHL015W-A
MTAFASLREPLVLANLKIKVHIYRMKR*
>YBR072C-A
MHILTRSSKNAFPRSRSRQDIHISSHIHRDTSNSALLKILVITRTRLDSFVKT*
>YGR204C-A
MQWNAFSFVSYVYLRYFISFRPNIVLASVRLSWYSII*
>YMR105W-A
MILVHTGNVLYPRFIVVAFTFEQRQGGRCKGGKATCMASVQSYKVTMQISSMTIIYPLFIFFSL*
>YLR285C-A
MFLRFYNRLLYIYNELFNFLRFTVFLPFCEFVPHHDLFYLIVTYQRTLVYKHKNDC*
>YLR159W
MKFQYALAKEQLGSNSRSGVKKLISKHHWLPEYYFSDLSFSVVQQWDSRAIEKTTIISCMRPANQEIYPLRHCETLRSQPCSLFSSLYARSFQSSCTLHVAEPSPGFHMYGCHT*
>YMR175W-A
MNCLCLCSLYSKSISAYFSEFSSTNIYKSYLRLPSVLYYVCMMHTMMPNQLDAVGIQSSESLLM*
>YOR011W-A
MAKSVFNFFHFEILEYLNRFVYHSQYFLPYYCSLEVLGKSRKNWTFQYWCLYITTDKKIIKKKDFYHR*
```

```
>YDR169C-A
MMRMSLKQMQSQRFSATSRVRMLLIGASGLRSSSSKLECPRFSNKHGRN*
>YJR151W-A
MLSLIFYLRFPSYIRG*
>YPL152W-A
MMVHLTLKSHLIKEELLWHALSPSYSCRYHGR*
>YPR159C-A
MATLDFTTKPLALVIYMSVVLLLMVGVPLLFSS*
>YLR161W
MKFQYALAKEQLGSNSRSGVKKLISKHHWLPEYYFSDLSFSVVQQWDSRAIEKTTIISCMRPANQEIYPLRHCETLRSQPCSLFSSLYARSFQSSCTLHVAEPSPGFHMYGCHT*
>YBL100W-C
MSRSIFFFSSLFLSPLNKIRNIGGKVENPWKSQIRDWKN*
>YDR246W-A
MRRLYRHLASFFLLPSCPGNTIQSITSYPANALLRSFRHVSTETPVRNRVHNRDSQSCPFFPLMDD*
>YGL188C-A
MLTTQKCESREGKNDEIFELGESNSDKILLKHGKCNLFSERKPVNH*
>YHL048C-A
MHDIWVITTSPACFEILYKYCKQKGRARMGGLIVKIIRFNHASV*
>YBR196C-B
MWVVLSKEKILLKKAYYAKTILFSALVLRGVRGE*
>YKL096C-B
MKLFILDYEKKRTKIGKGMARRELKMMNKKPDLYTIIVSYFSIFSLFFF*
>YOL164W-A
MFAKLYRGLTYVTNKILSTLLSHHSIHYCINSFKPTTLAIIILLHPCCRCNVYIIRCIHH*
>YOR032W-A
MRRALFIAGQTYLWLNLTHLLLIFSWSSTMAFSQSRRLLTPTVPCPTLLGIDFLILVLRHFDEIFI*
>YCR095W-A
MTVLIKLGLRILHVYKGFFRKVILKYFFFSSEHTKVNKKSSMHAFLCKIYKR*
>YOR161C-C
MSYTRVDHPTGKMACHLRQILASPLFFANYVLHAAIHYPSSDIRGDIL*
>YER175W-A
MRLHKTFICFSQNKRGCRNILQENSRMIFENKILIMILRQGIFFNISVSTKISF*
>YLR406C-A
MIQKPILLSSFLFLYIRALLHSIHPYIRTSVHLYTKKITSYNFLGVPFK*
>YOR394C-A
MIVNNTHILTLPPHAVSTLTCILIWHRHTDATVYIISSYPTLTFHSMAHLSLHQY*
>YMR315W-A
MTERKLLQLLRRPFISLSLFTALRACPLRPKSLIA*
```

**Table S1.** List and sequence of orphans found in *S. cerevisae* in fasta format.

```
>NP_652358.1
MPPPPPHHHHNPPPHGPPPHGPPHHGPPHHEPQPPFHTPVHPIYHEPPHGNSSGYNPPPPHGPYPYGPGYVAPGPPMHPR
>NP_001285519.1
MRVIFLIVVLAMVALAAVQGRRPPGLPQGGLGPLQGSGGLQPPQGIPQGPPQGLPQGPPQGGNSSSSEDSSSQESSSESSSEASA
>NP_001334703.1
MRTQVLSFCLAILLIVCLL
>NP_523436.1
MRTLILVTLVALVAVASAQGPGPWGPGGPGGPGGPGRGRGGPGRGPGGPGGPGGRGPGGPGGPGGPGGPGGPGGPGGPGCPGGPGGPGGPKPWGPPSNQTTSTTTEASTSTSTTTASSTTVSSTTESSTESSTESSTASSTE
>NP_608451.1
MRVIFLIVVLAMVALAAVQGRRPPGLPQGGLGPLQGSGGLQPPQGIPQGPPQGLPQGPPQGGNSSSSEDSSSQESSSESSSEASA
>NP_001286780.1
MRFQFVLAFGLIAILATAYAGGDTTGTGSTGTDTTGTGSTGTGSTGTGSTGSTGTGSTGTGSTGSTGTGSTGTGSTGSTGTGSTGSTATGSTGSTGSTTSGSPSTSTGSSSTNPSPSTTSPSTSPTTSTSPSDSSSSGSSSSSSSSKRNRKERRRRYRRRQRRQERRRERRERQRQQRRRG
>NP_001097828.1
MKYLSLSFLLCCLVAGVLSAPQFGYGFAPYGGYGGYGGYGGSYASASASSSAGGYQGFGYPGYGGYGGYGGFGGGYGGGYGQQYNQFSNYNNYGSSGYYGGYPFGR
>NP_001259864.1
MPPPPPHHHHNPPPHGPPPHGPPHHGPPHHEPQPPFHTPVHPIYHEPPHGNSSGYNPPPPHGPYPYGPGYVAPGPPMHPR
>NP_001334727.1
MLRYRLVSTHFLKQP
>NP_652283.1
MRFALLAVLLIGVIFAFVSAGGGGGGGGGQGGWQKNGGGGGGGGQGGWQKGGGGGGGGKHGGGGGGGGKHGGGNGGGGKHGGGGGGGGGGGKHGGGGW
>NP_608541.1
MRSFILIALFALIALAAAQGGPQGGQQGGQGGSQNGQQGGQQGGSQEGSQEGSQEGQQGGQQGGQQQGGPGGPWGLPPNGTELSNSTTTSTTEATSTESSSTEVSSS
>NP_001014551.1
MRAKWRKKRMRRLKRKRRKMRARSK
>NP_001334678.1
MLVWNDKQDTNFSFI
>NP_001286103.1
MCVRLVGSALSVANWASSACASSPVPRSVFSP
>NP_001303572.1
MAPPQDQPEDDIITKIPSIIGFLNLD
>NP_001138052.1
MSHDLDPTGTY
>NP_001162981.1
MDLKCCLLLLSLLFAGALARPEPDTPRPTLPPCSPWCAGQPPPPPGAVN
>NP_001162979.1
MKFLILFIALFAAAAALPQFGFGGGFGDQQQQQQGGFGGGFGDQQQQQGGFGGGFGGGFGDQQQQQQGGFGGGFGGGFGDQQQQQQQGW
>NP_001263026.1
MKLLILFLAALGAASARIRGHTIKG
>NP_001259862.1
MPPPPHQPHQPHPPKGPAHPGDHQPGPHPPHHGDPKPHHGHH
>NP_001162978.1
MKFLILFIALFAAAAALPQFGFGGGFGDQQQQQQGGFGGGFGDQQQQQGGFGGGFGGGFGDQQQQQQGGFGGGFGGGFGDQQQQQQQGW
>NP_001260648.1
MNSARHHRIGVFLSP
>NP_001259863.1
MPPPPHHDHHHHHGPPHHGPPHHGPPHHGPPHHGPPHHGPPHHGPPHHGPPHHDHHHHGPPHHGPPMHPR
>NP_001097701.1
MCCGPCGSCCGYYCCGPCCGPCGPRCGPCGSCCGPCGPCGPCCGPFGSCCGGCWC
>NP_001163495.1
MKLLSAALVLLMSSALAMAQKNTNTNENNIVIGKV
>NP_001334742.1
MDCCELPEIFLSFLNDLFFSDITDAEYLGDYYDNYN
>NP_001334726.1
MLRYRLVSTHFLKQP
>NP_001286449.1
MKLLIVVLALVAAVLARPQFGPGGQGGFGGQQGGFGGQGGFGGGPGFGGQGGFGGGPGEYGGQGGFGGGPGGYGGQGGFGGGPGGFGGQGGFGGGQGGFGEHHRHHGGHHGRF
>NP_001189120.1
MAHIGTFIAIALLAAVAQPACPGGCSGWVQHG
>NP_001245834.1
MPPPPHQPHQPHPPKGPAHPGDHQPGPHPPHHGDPKPHHGHH
>NP_001163438.1
MKSLLFFLLVILCLVGMAPARRKKREVEVWVRPSQNSYNEPCYYQGCQQ
>NP_001097010.1
MQTTITTRRLGLLGLLVVLGCIVAQGKPHGWSSGGGGGGWSSGGSGGSKVIISAWPSSGGGGWSSGGGSGGWKSGGDGGGGWSSGGSGGGWKSGGGSSGWKSGGGSGGWSSGGSSGWPSSG
SSGWKSGGSSGLSSALSSLSSWKSQALGSLSSGWKSGGGGGGWSSGGSSGGGWKSGGGSGGWSSGGSSAWPSKISSGWSSGGGGGGWSPGGGGGSGWSW
>NP_001245835.1
MPPPPHHDHHHHHGPPHHGPPHHGPPHHGPPHHGPPHHGPPHHGPPHHGPPHHDHHHHGPPHHGPPMHPR
>NP_001286239.1
MSWATTKQRHKCFNCSSGSNKNNRSSGSSSNNNNCAPSNKIISTSDFL
>NP_609439.1
MRLFIVLSVCVALASAGGYGGGGGGGGGHAGASASSSASAGAIGGGDGGGKLGGGYGSGGHGSGGLGSGGFGSGGDLGLGGGGVGGGPFAGGHAGGGVISGGGHSGGGGGFGGGPGFGSG
GHSGGGIGDGPGFGSGGYSGGGGGIGGGGGHSGGGSGIGGGPGFGGGIGGGGGHSGGGGGIGGGPGFGGGIGGSGSSASSSVGVIGGGHGGGGHGGGGFGPGGGGGGGFGPGGGGFGPGIG
GGGGGFGPGIGGGSGGGHFGGGLSHKHKGHGGGGGFKGGYGGGGGGGGGGGGGYGIGPAVSLGGGPGGGGSYGGGYGGGSGASASASASASAGAGGGYHG
>NP_001162976.1
MKFLILFIALFAAAAALPQFGFGGGFGDQQQQQQGGFGGGFGDQQQQQGGFGGGFGGGFGDQQQQQQGGFGGGFGGGFGDQQQQQQQGW
>NP_001189127.1
MKLLLPLTLAVIAICCMSFIECASSGDTATKTGTGTGTGTGTGTGTGTGTATTSTTVAPTTTSTTTAEATVHHRRIHRRRRRNLRRLRHRRAEAERRRRLG
>NP_723882.1
MKFLILFIALFAAAAALPQFGFGGGFGDQQQQQQGGFGGGFGDQQQQQGGFGGGFGGGFGDQQQQQQGGFGGGFGGGFGDQQQQQQQGW
>NP_729667.1
MASRIRREVTMRPILVLSLVILATLVVLSSQATSTSPTSSSSTSPTSSSSTSPTSSSSSTSSATTTTTTTTTTAATTTSTTTEKSKKKRNRRRRRRIIIRRRERRDERGERSRRERGDERGE
RSRRNRGSDGERIVRRYVVRERRFRRY
>NP_524254.1
MCCGPCGSCCGYYCCGPCCGPCGPRCGPCGSCCGPCGPCGPCCGPFGSCCGGCWC
>NP_611030.1
MKLLIVVLALVAAVLARPQFGPGGQGGFGGQQGGFGGQGGFGGGPGFGGQGGFGGGPGEYGGQGGFGGGPGGYGGQGGFGGGPGGFGGQGGFGGGQGGFGEHHRHHGGHHGRF
>NP_648438.1
MRPILALSLAVLATLVVLSSQATSTSPTSSSSTSPTSSSSTSPTSSSSTSPTSSSSSSDTTATTTTTTAATTTTTSTTEESKKKHRRRHRRRRIIIIRRVGGGLRRRRERDGDEDGGRVVRRIRVREGRFRRRDAFF
>NP_649406.2
MRFIFVLLLALLGCLLFAQQGCEATGTSTESSSDSTSASDTSTTASSSSDTTEASTSSDTTTVASSATTTTTSSSSSSSSSSSAAARRRRAAARRRRLARQRRRRQQRQRRQQRQRRRRQQQRRRRQQRQRSG
```

```
>NP_611823.2
MRFQFVLAFGLIAILATAYAGGDTTGTGSTGTDTTGTGSTGTGSTGTGSTGSTGTGSTGTGSTGSTGTGSTGTGSTGSTGTGSTGSTATGSTGSTGSTTSGSPSTSTGSSSTNPSPSTTSPS
TSPTTSTSPSDSSSSGSSSSSSSSKRNRKERRRRYRRRQRRQERRRERRERQRQQRRRG
>NP_001261798.1
MKSLLFFLLVILCLVGMAPARRKKREVEVWVRPSQNSYNEPCYYQGCQQ
>NP_001286240.1
MSWATTKQRHKCFNCSSGSNKNNRSSGSSSNNNNCAPSNKIISTSDFL
>NP_730309.1
MRFLLALSLAVLVVLILHSSQVSSTSTTTSTSASATTTTAATTTTDSSTTTTTESSTTTTASTKKKHRRRRVIIRRVIIRRGREGRSESGDNRRRVVVRVRRTAN
>NP_001287026.1
MKLVAITLVAFVAICLYSVDCTTSGTGSAATSGSGSSSASGSGSSSGSGTGTGTGTGTGTGTSTTLAPTTTTTTTTTVAPVKKVHRKRRHVIRKIVRRRAGERRRRG
>NP_730308.1
MRFLLALSLAVLVVLILHSSQVSSTSTTTSTSASATTTTAATTTTDSSTTTTTESSTTTTASTKKKHRRRRVIIRRVIIRRGREGRSESGDNRRRVVVRVRRTAN
>NP_729668.1
MKLVAITLVAFVAICLYSVDCTTSGTGSAATSGSGSSSASGSGSSSGSGTGTGTGTGTGTGTSTTLAPTTTTTTTTTVAPVKKVHRKRRHVIRKIVRRRAGERRRRG
>NP_001138051.1
MAAYLDPTGQY
>NP_001138050.1
MAAYLDPTGQY
>NP_001260533.1
MFWDLLGGIFEFLFGLLFPSSGSGDEPDSEEVETT
>NP_001334700.1
MKLIDVIIYLIAILIFIYLLLE
```

**Table S2.** List and sequence of orphans found in *D. Melanogaster* in fasta format.

```
>XP_015041786.1
EHEEEHEDEHEEEHEEEHEPKNTASDASCEEPSQDTVGGFGGAVVTLH
>XP_002132370.1
MKYFLSIIFGALSLTSAQYFGEDGIYGAAPNGPYGQQRPQQQQQAQTQDYFNGGFGYGYGPAFAAGLNQLQMQQQQQQLQQQQLDPYFYGGGPYGG
>XP_015042143.1
MHETNMSPVPNQDEGSQSCPQVDTLDQKEEPDQDSMKVVELGDELGQEEPLDYNDTSVEELVEKWKQRSEERDCQYASIGLQSEKILAELCSISEQIKQIQMRSNEEAADNQA
AGDQRDGNQGEIDQGNGDQENTEPQDDENKENSNRVDADQGYYQWDADQFFVYQGDAGQGSAGQGDSNQDDADTAPSSVSPAYFTF
>XP_015040782.1
MNFPSLLLLALVAGIAASHRIAQPPAAPEESHSTTTLQPPPPTSTTSTTTNAPAIATSTTCFTTTCTTPTTACNTTPCTTRTTSFSTTTTCSTTPYTTTTAPYTTTTTCGTTPYTTTKPCTTTPYTTTTTPSASLIEKLIPKRIG
>XP_015042265.1
MLFIQTGVCVSCGTMQQCETGQQRETKQKQKQQQQQRTPYTSSSSSPTSPHPQPPSEDCNANASANASDHAYAYASVC
>XP_015042264.1
MDNKQNHIAKLVRTSPRRVLRENKPCRKKVDFVPSCGTMQQCETGQQRETKQKQKQQQQQRTPYTSSSSSPTSPHPQPPSEDCNANASANASDHAYAYASVC
>XP_015042651.1
MCFCTVLVLKISVVLSILLLVLFNVLDIIEGVQNLNEPSSVRAYMRRYARISSLCKCLFSHW
>XP_015044418.1
MSAVFQFSLFSILLISISIWSSLIASLWAEEIDYAELKEKTLCIDDGMGHMMCY
>XP_015038371.1
MSNPLEESNRRVIAGGSGTGGDGGHGGKGGGGGGGGGGRGGKGGDGGGSNKESKDDPKAKDDKKSQEKENDQRRGKDGKDGEDGGSGSGGKGGDGGGAKGGKGGEGGGNQKESKDDPKTKEDGKKSQDGKDGKDGGSGRGGKGGDGGSGGGGGGRDGGKGGD
>XP_015036957.1
MAGTATATEMATVTAYQSQSQRPLATSPMSMTATATATAKATATMTVIEICGKQIKRVHSQR
>XP_015035445.1
MKFFIIFFALIVAVLAAGPGNHGEHGQQGHGQQAGSHGGQGGQQGFGHGGQGGQQGFGHGQQGNAQGHNKAGHGGH
>XP_015038770.1
MSNPLEESNRRVIAGGSGTGGDGGHGGKGGGGGGGGGGRGGKGGDGGGSNKESKDDPKAKDDKKSQEKENDQRRGKDGKDGEDGGSGSGGKGGDGGGAKGGKGGEGGGNQKESKDDPKTKEDGKKSQDGKDGKDGGSGRGGKGGDGGSGGGGGGRDGGKGGD
>XP_015039586.1
MKMKGHNYAYVLRFKCLASTIKTVYHPQQRQQHERQSQHQQRQHQRQQRLRQLKQQQNQQQQLRRQHQEQTR
>XP_015043478.1
SLPLSLYLSLSLSVCLSLGSSEASGVPDFMAVVQQFACNLSQQQQQQQQEQQQHAPSGGSLRLQLQLNS
>XP_002133927.1
MRSALLVLFLCGMLFAFASAGLLGGGGGGGGGYGGGGQGGWQNNGGGGGGHGGGGQGGWQKNGGGGGGQGGWQNNGGGGGGAQGGWQKNGGGGGGQGGWQNGGGGQGGWQNGGGSQGGWQNGGGSGGHGGGKSLAGNRGSSVSWQGGNGGGGGKHGGGGGGAGGKHGGGW
>XP_015042144.1
MSPVPNQDEGSQSCPQVDTLDQKEEPDQDSMKVVELGDELGQEEPLDYNDTSVEELVEKWKQRSEERDCQYASIGLQSEKILAELCSISEQIKQIQMRSNEEAADNQAAGDQRDGNQGEIDQGNGDQENTEPQDDENKENSNRVDADQGYYQWDADQFFVYQGDAGQGSAGQGDSNQDDADTAPSSVSPAYFTF
>XP_015042145.1
MSPVPNQDEGSQSCPQVDTLDQKEEPDQDSMKVVELGDELGQEEPLDYNDTSVEELVEKWKQRSEERDCQYASIGLQSEKILAELCSISEQIKQIQMRSNEEAADNQAAGDQRDGNQGEIDQGNGDQENTEPQDDENKENSNRVDADQGYYQWDADQFFVYQGDAGQGSAGQGDSNQDDADTAPSSVSPAYFTF
>XP_015034972.1
MKCHSFCSLFNIGTFRDFIFG
>XP_015040408.1
MSLAALSLILMLVFSISPLLSLEGKRMGP
>XP_002137838.2
MSAPQWGLGSGFGYGFAPYSGYGGSYASSSASAGASSSTGYQQGFGSPYGGYGGYGGYGGFGGFGGYGGYRQQYNQFSNQQNSYGSSGYGGFGYPFGYGR
>XP_015042623.1
MPQRIQTINMNPWSYLSLSLHAFPTRLFLQLLLIPTIAI
>XP_015042146.1
MSPVPNQDEGSQSCPQVDTLDQKEEPDQDSMKVVELGDELGQEEPLDYNDTSVEELVEKWKQRSEERDCQYASIGLQSEKILAELCSISEQIKQIQMRSNEEAADNQAAGDQRDGNQGEIDQGNGDQENTEPQDDENKENSNRVDADQGYYQWDADQFFVYQGDAGQGSAGQGDSNQDDADTAPSSVSPAYFTF
>XP_002137530.2
MKLFVCVLALAAILATTNAGLLSLLKGGGGSGGGYSGGSGYGGGSGYGGGYGAGYGGGYNSGYGGGHSRVDVYKVITTTHGGGHSGGYGGGYGGGYSGGYGGGYAANYGSSYGGGYGSQSGWWKKK
>XP_002133789.1
MLHFFNFNYFFGIGQQLLDVIGIYMMKWFGKRECHMHIVQTQTEAQPQVEAQTQTEHHYWTDDEAPEASQSDVDGSK
>XP_015042266.1
MQQCETGQQRETKQKQKQQQQQRTPYTSSSSSPTSPHPQPPSEDCNANASANASDHAYAYASVC
>XP_002135331.1
MKLSVLLWISLALLMSMAAAQGPAEAAQAGIGAAAGAGAQIGAQAGGGADAAEGMGVGMDAGVGGGAGAGAGAGAGAKGGAGVGGGGGGGAKGRR
>XP_015035443.1
MNFLIVLFALVAAAFAVPQYYGYSVGYPTYYGAEYPSYYGGEYPSYYGDRYSSYPDDSQESYESEEAQQEC
>XP_015042806.1
MPICISICEEPQLMSQQQRQQSGKAAKRQKQQPIGQIRVALSLRRREERQIEIERDR
>XP_015036675.1
MPVCLYLCQCQGMKKEYAPQEQALPQSQPQPQQQPQPHHPQPHHPQPQPQEQQQQHNGNGNSVNVEQRTSLLELAMIS
>XP_015041911.1
MQHDDAGRRLRTENKKKITSIEAWKSTEEKLQQPQQQQQQKGGWGGREGEVQWQFVFCFCFCFCFCTSSCFSLRLWLFLFMALSIPLCAFRLFLLFFLSCFLFLFHSFALLGLLCSTASTSHCYSRRSSASNYYF
SRSFSASMSYSSFSSAHHSYSKGSSASHSYSSLSSDSNSYSSRSSAFHSISFVLLVFIPILDVLLIQTPIPVVLLLLPMAIPVVLLLPIPIPVVLLMLIPLRDVFLLPIPIPVVLLLQISITVLLLWPINIPVVL
LLWVSSLFIRCFFFFNAFLFFSFLYFFNILLFPPFSI
>XP_015042263.1
MDNKQNHIAKLVRTSPRRVLRENKPCRKKVDFVPSSCGTMQQCETGQQRETKQKQKQQQQQRTPYTSSSSSPTSPHPQPPSEDCNANASANASDHAYAYASVC
>XP_002133104.2
MKILIVLAVLVTVASALPQFGFGGPGFGGPGFGGPGFGGGGYGRPGFGGGPGFGGGFGGGPGYGGFGGGPGYGGFGGSPASAAGGGAASSSASSSASASGGGGFYG
>XP_015036494.1
MGQKQGQKQGQKQGQNQGQNQGPEPEPVTRKYETISKPSWIASHSTIENMSLENGKYERSTGIDWRTTRDSATAEIHSN
>XP_015041143.1
MSSICQRSLDVPVRVAARARNEQQTATNVISGAQTLWKRQNPHKSQSVAAAAPAAAAAGPEPEAQKPKARRRSACRRHEGWMTRTRTTTTMTLALAGALFWWWRKQEEGA
>XP_015042653.1
MCFCTVLVLKISVVLSILLLVLFNVLDIIEGVQNLNEPSSVRAYMRRYARISSLCK
>XP_015041370.1
MLRIGVRVGVGVYNFQSWNPVGGEVGLRLAGGMSHLDMDRDLDLDLDLTIYSWRRQIWN
>XP_002134556.1
MQQCETGQQRETKQKQKQQQQQRTPYTSSSSSPTSPHPQPPSEDCNANASANASDHAYAYASVC
>XP_002134480.1
MSPVPNQDEGSQSCPQVDTLDQKEEPDQDSMKVVELGDELGQEEPLDYNDTSVEELVEKWKQRSEERDCQYASIGLQSEKILAELCSISEQIKQIQMRSNEEAADNQAAGDQRDGNQGEIDQGNGDQENTEPQDDEN
KENSNRVDADQGYYQWDADQFFVYQGDAGQGSAGQGDSNQDDADTAPSSVSPAYFTF
```

**Table S3.** List and sequence of orphans found in *D. pseudoobscura* in fasta format.

| query_id | RNA_seq_exp_1 | RNA_seq_exp_2 | RNA_seq_exp_3 | RNA_seq_exp_4 | RNA_seq_exp_5 | RNA_seq_exp_6 | RNA_seq_exp_7 |
|---|---|---|---|---|---|---|---|
| YER175W-A | 3.25 | 3.18 | 4.06 | 1.59 | 4.02 | 4.92 | 5.52 |
| YAL037C-A | 11.40 | 1.21 | 0.72 | 2.84 | 0.00 | 0.68 | 0.55 |
| YBR196C-B | 2.17 | 17.18 | 16.00 | 13.57 | 10.56 | 12.55 | 15.46 |
| YOL164W-A | 3.17 | 2.45 | 1.83 | 2.87 | 0.00 | 0.00 | 0.28 |
| YDR114C | 1.58 | 1.73 | 0.22 | 0.00 | 0.48 | 0.82 | 0.33 |
| YIR018C-A | 2.17 | 1.90 | 4.37 | 2.29 | 1.07 | 1.36 | 3.30 |
| YGR121W-A | 1.00 | 13.33 | 10.53 | 8.27 | 8.17 | 10.40 | 11.69 |
| YDR246W-A | 1.00 | 9.87 | 15.98 | 8.36 | 12.44 | 9.63 | 12.06 |
| YOR161C-C | 1.00 | 3.57 | 4.56 | 2.86 | 3.01 | 0.43 | 3.10 |
| YOR011W-A | 1.00 | 0.00 | 0.97 | 0.00 | 0.00 | 0.00 | 0.24 |
| YMR230W-A | 0.00 | 17.62 | 16.29 | 10.56 | 10.12 | 11.90 | 11.23 |
| YKL096C-B | 2.81 | 1.50 | 0.89 | 0.70 | 2.46 | 2.08 | 0.67 |
| YAL064W | 0.00 | 3.15 | 4.45 | 3.13 | 2.06 | 2.84 | 3.54 |
| YJL136W-A | 1.00 | 4.93 | 4.01 | 5.67 | 0.88 | 2.25 | 0.00 |
| YOR032W-A | 0.00 | 4.65 | 3.33 | 5.23 | 3.29 | 4.66 | 4.77 |
| YMR315W-A | 0.00 | 4.18 | 5.60 | 5.86 | 2.74 | 4.65 | 7.52 |
| YLR157W-D | 0.00 | 0.00 | 0.00 | 0.25 | 0.00 | 0.00 | 0.00 |
| YDR169C-A | 0.00 | 3.00 | 2.23 | 1.75 | 0.98 | 0.83 | 1.69 |
| YOR381W-A | 1.00 | 2.90 | 3.19 | 2.19 | 2.19 | 1.49 | 1.81 |
| YHL048C-A | 0.00 | 3.61 | 6.96 | 2.73 | 1.09 | 1.39 | 0.38 |
| YMR105W-A | 0.00 | 2.49 | 3.09 | 0.81 | 1.13 | 2.56 | 5.18 |
| YOR008C-A | 0.00 | 2.49 | 1.19 | 1.87 | 1.31 | 1.66 | 1.57 |
| YOR394C-A | 0.00 | 3.57 | 1.20 | 3.13 | 0.44 | 2.23 | 1.20 |
| YGL188C-A | 0.00 | 13.83 | 9.99 | 12.32 | 12.02 | 22.19 | 11.85 |
| YLR285C-A | 0.00 | 362.65 | 323.47 | 269.62 | 360.76 | 401.65 | 339.69 |
| YOL155W-A | 0.00 | 0.83 | 0.50 | 0.00 | 0.55 | 0.00 | 0.00 |
| YMR175W-A | 0.00 | 1.34 | 0.34 | 1.89 | 0.00 | 0.96 | 0.52 |
| YIL046W-A | 0.00 | 2.72 | 3.25 | 2.55 | 3.57 | 0.76 | 1.84 |
| YOR293C-A | 4.49 | 0.50 | 0.00 | 0.00 | 0.00 | 0.42 | 0.00 |
| YLR412C-A | 0.00 | 0.36 | 0.97 | 1.52 | 0.71 | 2.11 | 0.98 |
| YDR194W-A | 0.00 | 0.24 | 0.44 | 0.00 | 0.00 | 0.00 | 0.00 |
| YMR247W-A | 2.81 | 0.00 | 0.00 | 0.00 | 0.51 | 0.00 | 0.00 |
| YNL067W-B | 0.00 | 0.53 | 0.95 | 1.12 | 0.00 | 0.00 | 0.00 |
| YAL063C-A | 0.00 | 5.39 | 7.35 | 4.15 | 4.29 | 3.00 | 4.33 |
| YGR204C-A | 1.00 | 2.97 | 1.18 | 3.70 | 1.30 | 2.20 | 3.11 |
| YCR095W-A | 2.32 | 8.25 | 10.53 | 12.90 | 8.80 | 9.04 | 11.77 |
| YPL152W-A | 0.00 | 1.52 | 2.04 | 3.73 | 2.99 | 0.63 | 2.05 |
| YNL277W-A | 1.00 | 3.17 | 2.12 | 4.17 | 0.78 | 0.66 | 1.60 |
| YGL006W-A | 0.00 | 0.00 | 0.00 | 0.00 | 0.00 | 0.00 | 0.46 |
| YGR174W-A | 4.46 | 39.39 | 24.77 | 35.86 | 29.78 | 26.01 | 54.35 |
| YOR376W-A | 2.58 | 52.84 | 49.82 | 60.36 | 60.91 | 52.91 | 92.09 |
| YMR272W-B | 0.00 | 0.35 | 0.00 | 0.49 | 0.00 | 0.00 | 0.00 |
| YBR072C-A | 2.32 | 1.39 | 1.65 | 1.62 | 0.00 | 0.39 | 0.62 |
| YBR296C-A | 0.00 | 1.25 | 3.92 | 0.44 | 1.23 | 1.57 | 2.53 |
| YLR406C-A | 0.00 | 1.75 | 2.68 | 1.75 | 0.49 | 0.83 | 0.67 |
| YOL038C-A | 3.46 | 31.74 | 30.13 | 33.01 | 15.41 | 30.74 | 30.68 |
| YBL100W-C | 1.00 | 1.56 | 0.00 | 0.88 | 1.23 | 0.00 | 1.69 |
| YHL015W-A | 0.00 | 1.79 | 1.60 | 1.89 | 0.88 | 0.00 | 0.61 |
| YPL038W-A | 0.00 | 13.84 | 16.03 | 9.58 | 17.24 | 13.34 | 14.74 |
| YJL077W-A | 0.00 | 167.96 | 154.82 | 169.56 | 84.25 | 81.63 | 85.91 |
| YBR182C-A | 1.00 | 2.11 | 1.03 | 2.42 | 1.13 | 1.92 | 1.04 |
| YHR022C-A | 0.00 | 0.84 | 0.75 | 0.59 | 0.00 | 1.40 | 0.00 |

**Table S4.** For the 52 *S. cerevisiae* genes out of the 63 orphans found by blastp+tblastn, it is shown the expression level in 7 different RNA-Seq experiments. The values in the columns corresponds to the RNA expression data from the two experiments taken into consideration. The values in the first column (RNA_seq_exp_1) are the log2 tag count reported in [1]. The values in the columns RNA_seq_exp_2, RNA_seq_exp_3 and RNA_seq_exp_4 are the Reads Per Kilobase Million (RPKM) values for the three replicates of the Disome 10 (chromosome X disomy) reported in [2]. The values in the columns RNA_seq_exp_5, RNA_seq_exp_6 and RNA_seq_exp_7 are the RPKM values for the three replicates of the wild type haploid reported in [2].

| Experiment | Name RNA-Seq dataset from modENCODE |
|---|---|
| exp1 | PolyA-RNA-Developmental-Stage-Adult-Male-Strain-Y-cn-bw-sp-RNA-seq-Rep-1-Dmel_r5-32-modENCODE_2027 |
| exp2 | PolyA-RNA-Developmental-Stage-Embryos-0-4-hr-Strain-Y-cn-bw-sp-RNA-seq-Rep-1-Dmel_r5-32-modENCODE_2010 |
| exp3 | PolyA-RNA-Developmental-Stage-Embryos-16-20-hr-Strain-Y-cn-bw-sp-RNA-seq-Rep-1-Dmel_r5-32-modENCODE_2022 |
| exp4 | PolyA-RNA-Developmental-Stage-Embryos-20-24-hr-Strain-Y-cn-bw-sp-RNA-seq-Rep-1-Dmel_r5-32-modENCODE_2023 |
| exp5 | PolyA-RNA-Developmental-Stage-Embryos-4-8-hr-Strain-Y-cn-bw-sp-RNA-seq-Rep-1-Dmel_r5-32-modENCODE_2019 |
| exp6 | PolyA-RNA-Developmental-Stage-Embryos-8-12-hr-Strain-Y-cn-bw-sp-RNA-seq-Rep-1-Dmel_r5-32-modENCODE_2020 |
| exp7 | PolyA-RNA-Developmental-Stage-Larvae-L2-stage-Strain-Y-cn-bw-sp-RNA-seq-Rep-1-Dmel_r5-32-modENCODE_2025 |
| exp8 | PolyA-RNA-Developmental-Stage-Larvae-L3-stage-Strain-Y-cn-bw-sp-RNA-seq-Rep-1-Dmel_r5-32-modENCODE_2029 |
| exp9 | PolyA-RNA-Developmental-Stage-Pupae-Strain-Y-cn-bw-sp-RNA-seq-Rep-1-Dmel_r5-32-modENCODE_2030 |
| exp10 | total-RNA-Developmental-Stage-Adult-Female-8-days-Strain-Oregon-R-Tissue-Dmel-Female-heads-RNA-seq-Rep-1-Dmel_r5-32-modENCODE_3083-33 |
| exp11 | total-RNA-Developmental-Stage-Adult-Female-8-days-Strain-Oregon-R-Tissue-Dmel-Female-heads-RNA-seq-Rep-1-Dmel_r5-32-modENCODE_3083-34 |
| exp12 | total-RNA-Developmental-Stage-Adult-Female-8-days-Strain-Oregon-R-Tissue-Dmel-Female-heads-RNA-seq-Rep-1-Dmel_r5-32-modENCODE_3083-35 |
| exp13 | total-RNA-Developmental-Stage-Adult-Female-8-days-Strain-Oregon-R-Tissue-Dmel-Female-heads-RNA-seq-Rep-1-Dmel_r5-32-modENCODE_3083-39 |
| exp14 | total-RNA-Developmental-Stage-Adult-Female-8-days-Strain-Oregon-R-Tissue-Dmel-Female-heads-RNA-seq-Rep-1-Dmel_r5-32-modENCODE_3083-40 |
| exp15 | total-RNA-Developmental-Stage-Adult-Female-8-days-Strain-Oregon-R-Tissue-Dmel-Female-heads-RNA-seq-Rep-1-Dmel_r5-32-modENCODE_3083-41 |
| exp16 | total-RNA-Developmental-Stage-Adult-Female-8-days-Strain-Oregon-R-Tissue-Dmel-Female-heads-RNA-seq-Rep-1-Dmel_r5-32-modENCODE_3083-42 |
| exp17 | total-RNA-Developmental-Stage-Adult-Female-8-days-Strain-Oregon-R-Tissue-Dmel-Female-heads-RNA-seq-Rep-1-Dmel_r5-32-modENCODE_3083-43 |
| exp18 | total-RNA-Developmental-Stage-Adult-Female-8-days-Strain-Oregon-R-Tissue-Dmel-Female-heads-RNA-seq-Rep-1-Dmel_r5-32-modENCODE_3083-44 |
| exp19 | total-RNA-Developmental-Stage-Adult-Female-8-days-Strain-Oregon-R-Tissue-Dmel-Female-heads-RNA-seq-Rep-1-Dmel_r5-32-modENCODE_3083-45 |
| exp20 | total-RNA-Developmental-Stage-Adult-Female-Whole-Species-Strain-Dmel-y1w67c23-RNA-seq-Rep-1-Dmel_r5-32-modENCODE_3631-7 |
| exp21 | total-RNA-Developmental-Stage-Adult-Female-Whole-Species-Strain-Dmel-y1w67c23-RNA-seq-Rep-1-Dmel_r5-32-modENCODE_3631-8 |
| exp22 | total-RNA-Developmental-Stage-Adult-Male-8-days-Strain-Oregon-R-Tissue-Dmel-Male-heads-RNA-seq-Rep-1-Dmel_r5-32-modENCODE_3084-36 |
| exp23 | total-RNA-Developmental-Stage-Adult-Male-8-days-Strain-Oregon-R-Tissue-Dmel-Male-heads-RNA-seq-Rep-1-Dmel_r5-32-modENCODE_3084-37 |
| exp24 | total-RNA-Developmental-Stage-Adult-Male-8-days-Strain-Oregon-R-Tissue-Dmel-Male-heads-RNA-seq-Rep-1-Dmel_r5-32-modENCODE_3084-38 |
| exp25 | total-RNA-Developmental-Stage-Adult-Male-8-days-Strain-Oregon-R-Tissue-Dmel-Male-heads-RNA-seq-Rep-1-Dmel_r5-32-modENCODE_3084-46 |
| exp26 | total-RNA-Developmental-Stage-Adult-Male-8-days-Strain-Oregon-R-Tissue-Dmel-Male-heads-RNA-seq-Rep-1-Dmel_r5-32-modENCODE_3084-47 |
| exp27 | total-RNA-Developmental-Stage-Adult-Male-8-days-Strain-Oregon-R-Tissue-Dmel-Male-heads-RNA-seq-Rep-1-Dmel_r5-32-modENCODE_3084-48 |
| exp28 | total-RNA-Developmental-Stage-Adult-Male-8-days-Strain-Oregon-R-Tissue-Dmel-Male-heads-RNA-seq-Rep-1-Dmel_r5-32-modENCODE_3084-49 |
| exp29 | total-RNA-Developmental-Stage-Adult-Male-8-days-Strain-Oregon-R-Tissue-Dmel-Male-heads-RNA-seq-Rep-1-Dmel_r5-32-modENCODE_3084-50 |
| exp30 | total-RNA-Developmental-Stage-Adult-Male-8-days-Strain-Oregon-R-Tissue-Dmel-Male-heads-RNA-seq-Rep-1-Dmel_r5-32-modENCODE_3084-51 |
| exp31 | total-RNA-Developmental-Stage-Adult-Male-8-days-Strain-Oregon-R-Tissue-Dmel-Male-heads-RNA-seq-Rep-1-Dmel_r5-32-modENCODE_3084-52 |
| exp32 | total-RNA-Developmental-Stage-Adult-Male-Whole-Species-Strain-Dmel-y1w67c23-RNA-seq-Rep-1-Dmel_r5-32-modENCODE_3632-09 |
| exp33 | total-RNA-Developmental-Stage-Adult-Male-Whole-Species-Strain-Dmel-y1w67c23-RNA-seq-Rep-1-Dmel_r5-32-modENCODE_3632-10 |

**Table S5.** The table shows the name of the datasets downloaded from modENCODE

| queryid | exp1 | exp2 | exp3 | exp4 | exp5 | exp6 | exp7 | exp8 | exp9 | exp10 | exp11 | exp12 | exp13 | exp14 | exp15 | exp16 |
|---|---|---|---|---|---|---|---|---|---|---|---|---|---|---|---|---|
| NP_001259864.1 | 0.00 | 0.00 | 0.00 | 6.96 | 0.00 | 0.00 | 6.95 | 0.00 | 0.00 | 0.00 | 0.00 | 0.00 | 0.00 | 0.00 | 0.00 | 0.00 |
| NP_649406.2 | 0.00 | 4.72 | 232.24 | 2649.95 | 3.69 | 80.83 | 56.72 | 0.00 | 7.38 | 0.00 | 0.00 | 0.00 | 0.00 | 0.00 | 0.00 | 0.00 |
| NP_001259862.1 | 0.00 | 0.00 | 0.00 | 3.73 | 0.00 | 0.00 | 1.49 | 0.00 | 0.00 | 0.00 | 0.00 | 0.00 | 0.00 | 0.00 | 0.00 | 0.00 |
| NP_609439.1 | 8071.99 | 0.00 | 9.61 | 6.57 | 0.00 | 2.18 | 392.68 | 1242.86 | 66.38 | 30853.70 | 23350.64 | 21319.15 | 7628.59 | 5262.62 | 9626.64 | 7851.16 |
| NP_001189120.1 | 21098.78 | 0.00 | 0.00 | 11.59 | 0.00 | 0.00 | 0.00 | 0.00 | 0.00 | 0.00 | 0.00 | 0.00 | 0.00 | 0.00 | 0.00 | 0.00 |
| NP_001261798.1 | 71.03 | 0.00 | 0.00 | 0.00 | 0.00 | 0.00 | 0.00 | 0.00 | 0.00 | 0.00 | 0.00 | 0.00 | 0.00 | 0.00 | 0.00 | 0.00 |
| NP_730309.1 | 0.00 | 0.00 | 3.25 | 48.94 | 0.00 | 0.00 | 159.98 | 0.00 | 0.00 | 3.76 | 0.00 | 0.00 | 0.00 | 0.00 | 0.00 | 0.00 |
| NP_608541.1 | 386.85 | 0.00 | 2.97 | 357.44 | 0.00 | 0.00 | 34.08 | 0.00 | 4.43 | 0.00 | 0.00 | 0.00 | 0.00 | 0.00 | 0.00 | 0.00 |
| NP_001163495.1 | 2871.98 | 0.00 | 0.00 | 0.00 | 0.00 | 0.00 | 0.00 | 0.00 | 0.00 | 0.00 | 0.00 | 0.00 | 0.00 | 0.00 | 0.00 | 0.00 |
| NP_001263026.1 | 34515.83 | 0.00 | 0.00 | 0.00 | 0.00 | 0.00 | 0.00 | 0.00 | 0.00 | 0.00 | 0.00 | 0.00 | 0.00 | 0.00 | 0.00 | 0.00 |
| NP_730308.1 | 0.00 | 0.00 | 0.00 | 21.93 | 0.00 | 0.00 | 188.35 | 0.00 | 0.00 | 0.00 | 0.00 | 0.00 | 0.00 | 0.00 | 0.00 | 0.00 |
| NP_001097010.1 | 296.63 | 0.00 | 75.86 | 67.68 | 0.00 | 7.49 | 16.22 | 0.00 | 531.90 | 1.90 | 3.47 | 1.05 | 0.00 | 0.00 | 0.00 | 0.00 |
| NP_729667.1 | 0.00 | 0.00 | 3.09 | 0.00 | 0.00 | 0.00 | 202.47 | 415.91 | 0.00 | 0.00 | 0.00 | 0.00 | 0.00 | 0.00 | 0.00 | 0.00 |
| NP_001189127.1 | 0.00 | 0.00 | 13.71 | 229.76 | 0.00 | 0.00 | 929.20 | 10.99 | 0.00 | 0.00 | 0.00 | 0.00 | 0.00 | 0.00 | 0.00 | 0.00 |
| NP_652283.1 | 1132.02 | 0.00 | 2.60 | 3.56 | 0.00 | 0.00 | 0.00 | 0.00 | 70.00 | 219.50 | 211.62 | 214.10 | 116.52 | 98.55 | 118.30 | 128.86 |
| NP_001260533.1 | 19.78 | 0.00 | 0.00 | 0.00 | 0.00 | 0.00 | 9.38 | 0.00 | 12.81 | 0.00 | 0.00 | 5.47 | 0.00 | 40.58 | 0.00 | 0.00 |
| NP_648438.1 | 3.24 | 0.00 | 0.00 | 53.78 | 0.00 | 0.00 | 838.01 | 9.01 | 0.00 | 0.00 | 0.00 | 0.00 | 0.00 | 0.00 | 0.00 | 0.00 |
| NP_523436.1 | 3197.59 | 0.00 | 2.37 | 55.12 | 0.00 | 0.00 | 500.06 | 91.24 | 3.54 | 383.14 | 328.76 | 350.46 | 183.19 | 156.97 | 165.65 | 150.79 |
| NP_001285519.1 | 0.00 | 0.00 | 0.00 | 0.00 | 0.00 | 0.00 | 0.00 | 0.00 | 4.81 | 0.00 | 0.00 | 0.00 | 0.00 | 0.00 | 0.00 | 0.00 |
| NP_001014551.1 | 20125.50 | 62009.40 | 54113.43 | 83518.84 | 81016.68 | 83832.75 | 89363.38 | 38290.87 | 22559.17 | 47527.09 | 60348.97 | 61661.62 | 41099.11 | 37231.28 | 46088.52 | 43247.21 |
| NP_001097828.1 | 7.20 | 0.00 | 0.00 | 179.35 | 0.00 | 0.00 | 39.24 | 0.00 | 0.00 | 0.00 | 0.00 | 0.00 | 0.00 | 0.00 | 0.00 | 0.00 |
| NP_001162981.1 | 0.00 | 0.00 | 811.97 | 308.56 | 14.98 | 27.33 | 46.57 | 426.07 | 104.78 | 0.00 | 0.00 | 3.19 | 0.00 | 0.00 | 0.00 | 0.00 |
| NP_001286449.1 | 0.00 | 0.00 | 5.09 | 118.41 | 0.00 | 0.00 | 20.41 | 10.89 | 0.00 | 11.76 | 9.51 | 6.49 | 6.90 | 0.00 | 5.93 | 17.99 |
| NP_001303572.1 | 29.16 | 0.00 | 7.59 | 653.84 | 3.78 | 27.58 | 150.66 | 68.95 | 101.96 | 11.68 | 24.80 | 9.67 | 0.00 | 0.00 | 0.00 | 8.94 |
| NP_001287026.1 | 8.41 | 0.00 | 0.00 | 0.00 | 0.00 | 0.00 | 0.00 | 0.00 | 3.63 | 0.00 | 0.00 | 0.00 | 0.00 | 0.00 | 0.00 | 0.00 |

| queryid | exp17 | exp18 | exp19 | exp20 | exp21 | exp22 | exp23 | exp24 | exp25 | exp26 | exp27 | exp28 | exp29 | exp30 | exp31 | exp32 | exp33 |
|---|---|---|---|---|---|---|---|---|---|---|---|---|---|---|---|---|---|
| NP_001259864.1 | 0.00 | 0.00 | 0.00 | 0.0 | 0.00 | 0.00 | 0.00 | 0.00 | 0.00 | 0.00 | 0.00 | 0.00 | 0.00 | 0.00 | 0.00 | 0.00 | 0.0 |
| NP_649406.2 | 0.00 | 0.00 | 0.00 | 0.0 | 0.00 | 0.00 | 0.00 | 0.00 | 0.00 | 0.00 | 0.00 | 0.00 | 0.00 | 0.00 | 0.00 | 0.00 | 0.0 |
| NP_001259862.1 | 0.00 | 0.00 | 0.00 | 0.0 | 0.00 | 0.00 | 0.00 | 0.00 | 0.00 | 0.00 | 0.00 | 0.00 | 0.00 | 0.00 | 0.00 | 0.00 | 0.0 |
| NP_609439.1 | 10337.87 | 8138.57 | 27340.77 | 0.0 | 92.34 | 37864.73 | 28761.81 | 27872.69 | 23627.73 | 30105.23 | 29079.13 | 26303.58 | 26649.07 | 30009.77 | 40396.82 | 0.00 | 64.7 |
| NP_001189120.1 | 0.00 | 0.00 | 0.00 | 0.0 | 0.00 | 0.00 | 0.00 | 0.00 | 0.00 | 0.00 | 0.00 | 0.00 | 0.00 | 0.00 | 0.00 | 0.00 | 0.0 |
| NP_001261798.1 | 0.00 | 0.00 | 0.00 | 0.0 | 0.00 | 0.00 | 0.00 | 0.00 | 0.00 | 0.00 | 0.00 | 0.00 | 0.00 | 0.00 | 0.00 | 0.00 | 0.0 |
| NP_730309.1 | 0.00 | 0.00 | 0.00 | 0.0 | 0.00 | 0.00 | 4.49 | 4.46 | 0.00 | 0.00 | 4.83 | 0.00 | 0.00 | 0.00 | 0.00 | 0.00 | 0.0 |
| NP_608541.1 | 0.00 | 0.00 | 0.00 | 0.0 | 0.00 | 0.00 | 0.00 | 2.04 | 0.00 | 0.00 | 0.00 | 0.00 | 0.00 | 0.00 | 0.00 | 0.00 | 0.0 |
| NP_001163495.1 | 0.00 | 0.00 | 0.00 | 0.0 | 0.00 | 0.00 | 0.00 | 0.00 | 0.00 | 0.00 | 0.00 | 0.00 | 0.00 | 0.00 | 0.00 | 0.00 | 0.0 |
| NP_001263026.1 | 0.00 | 0.00 | 0.00 | 0.0 | 0.00 | 0.00 | 0.00 | 0.00 | 0.00 | 0.00 | 0.00 | 0.00 | 0.00 | 0.00 | 0.00 | 0.00 | 0.0 |
| NP_730308.1 | 0.00 | 0.00 | 0.00 | 0.0 | 0.00 | 0.00 | 0.00 | 0.00 | 7.46 | 0.00 | 0.00 | 0.00 | 0.00 | 0.00 | 0.00 | 0.00 | 0.0 |
| NP_001097010.1 | 0.00 | 0.00 | 2.18 | 0.0 | 0.00 | 0.00 | 1.14 | 1.13 | 0.00 | 4.92 | 0.00 | 2.43 | 0.00 | 0.00 | 3.72 | 0.00 | 0.0 |
| NP_729667.1 | 0.00 | 0.00 | 0.00 | 0.0 | 0.00 | 0.00 | 0.00 | 0.00 | 0.00 | 0.00 | 0.00 | 0.00 | 0.00 | 0.00 | 0.00 | 0.00 | 0.0 |
| NP_001189127.1 | 0.00 | 0.00 | 0.00 | 0.0 | 0.00 | 0.00 | 0.00 | 0.00 | 0.00 | 0.00 | 0.00 | 0.00 | 0.00 | 0.00 | 0.00 | 0.00 | 0.0 |
| NP_652283.1 | 152.38 | 116.83 | 220.44 | 0.0 | 0.00 | 45.27 | 59.28 | 58.91 | 67.89 | 50.51 | 42.52 | 26.91 | 45.42 | 42.70 | 91.11 | 0.00 | 0.0 |
| NP_001260533.1 | 0.00 | 0.00 | 0.00 | 0.0 | 0.00 | 0.00 | 5.92 | 0.00 | 15.98 | 12.80 | 12.73 | 0.00 | 0.00 | 11.72 | 19.37 | 0.00 | 0.0 |
| NP_648438.1 | 0.00 | 0.00 | 0.00 | 0.0 | 0.00 | 0.00 | 0.00 | 1.93 | 0.00 | 0.00 | 0.00 | 4.15 | 0.00 | 0.00 | 0.00 | 0.00 | 0.0 |
| NP_523436.1 | 238.86 | 194.95 | 253.93 | 0.0 | 1275.26 | 8.83 | 16.35 | 9.75 | 4.41 | 3.54 | 10.55 | 10.50 | 10.33 | 0.00 | 13.38 | 179.12 | 0.0 |
| NP_001285519.1 | 0.00 | 0.00 | 0.00 | 0.0 | 0.00 | 0.00 | 0.00 | 0.00 | 0.00 | 0.00 | 0.00 | 0.00 | 0.00 | 0.00 | 0.00 | 0.00 | 0.0 |
| NP_001014551.1 | 46312.45 | 44030.33 | 49003.51 | 0.0 | 1343.57 | 20141.22 | 25559.18 | 25853.76 | 21685.83 | 22860.04 | 22383.54 | 22763.26 | 22363.49 | 22018.43 | 21199.08 | 0.00 | 0.0 |
| NP_001097828.1 | 0.00 | 0.00 | 0.00 | 0.0 | 0.00 | 0.00 | 0.00 | 0.00 | 0.00 | 0.00 | 0.00 | 0.00 | 0.00 | 0.00 | 0.00 | 0.00 | 0.0 |
| NP_001162981.1 | 0.00 | 0.00 | 0.00 | 0.0 | 0.00 | 0.00 | 0.00 | 0.00 | 0.00 | 0.00 | 0.00 | 0.00 | 0.00 | 0.00 | 0.00 | 0.00 | 0.0 |
| NP_001286449.1 | 0.00 | 0.00 | 6.73 | 0.0 | 0.00 | 4.21 | 9.37 | 6.98 | 6.32 | 2.53 | 10.08 | 2.51 | 9.87 | 4.64 | 1.92 | 0.00 | 0.0 |
| NP_001303572.1 | 24.66 | 0.00 | 23.41 | 0.0 | 0.00 | 3.14 | 8.72 | 10.40 | 4.71 | 18.87 | 7.51 | 18.67 | 7.35 | 10.37 | 14.27 | 0.00 | 0.0 |
| NP_001287026.1 | 0.00 | 0.00 | 0.00 | 0.0 | 0.00 | 0.00 | 0.00 | 0.00 | 0.00 | 0.00 | 0.00 | 0.00 | 0.00 | 0.00 | 0.00 | 0.00 | 0.0 |

**Table S6.** For the 25 *D. melanogaster* genes out of the 54 orphans found by blastp+tblastn, it is shown the expression level in 33 different RNA-Seq datasets. The values in the columns corresponds to the RNA expression data in Reads Per Kilobase Million (RPKM).

| Experiment exp1 | Name RNA-Seq dataset from modENCODE |
|---|---|
| exp1 | total-RNA-Developmental-Stage-Adult-Female-Strain-D-pseudoobscura-wild-type-Tissue-Dpse-Female-heads-RNA-seq-Rep-1-Dpse_r2-4-modENCODE_2980-82 |
| exp2 | total-RNA-Developmental-Stage-Adult-Female-Strain-D-pseudoobscura-wild-type-Tissue-Dpse-Female-heads-RNA-seq-Rep-1-Dpse_r2-4-modENCODE_2980-83 |
| exp3 | total-RNA-Developmental-Stage-Adult-Female-Strain-D-pseudoobscura-wild-type-Tissue-Dpse-Female-heads-RNA-seq-Rep-1-Dpse_r2-4-modENCODE_2980-84 |
| exp4 | total-RNA-Developmental-Stage-Adult-Female-Strain-D-pseudoobscura-wild-type-Tissue-Dpse-Female-heads-RNA-seq-Rep-1-Dpse_r2-4-modENCODE_2980-85 |
| exp5 | total-RNA-Developmental-Stage-Adult-Female-Strain-D-pseudoobscura-wild-type-Tissue-Dpse-Female-heads-RNA-seq-Rep-1-Dpse_r2-4-modENCODE_2980-86 |
| exp6 | total-RNA-Developmental-Stage-Adult-Female-Strain-D-pseudoobscura-wild-type-Tissue-Dpse-Female-heads-RNA-seq-Rep-1-Dpse_r2-4-modENCODE_2980-87 |
| exp7 | total-RNA-Developmental-Stage-Adult-Female-Strain-D-pseudoobscura-wild-type-Tissue-Dpse-Female-heads-RNA-seq-Rep-1-Dpse_r2-4-modENCODE_2980-88 |
| exp8 | total-RNA-Developmental-Stage-Adult-Female-Strain-D-pseudoobscura-wild-type-Tissue-Dpse-Female-heads-RNA-seq-Rep-1-Dpse_r2-4-modENCODE_2980-89 |
| exp9 | total-RNA-Developmental-Stage-Adult-Female-Strain-D-pseudoobscura-wild-type-Tissue-Dpse-Female-heads-RNA-seq-Rep-1-Dpse_r2-4-modENCODE_2980-90 |
| exp10 | total-RNA- Developmental-Stage-Adult-Female-Strain-D-pseudoobscura-wild-type-Tissue-Dpse-Female-heads-RNA-seq-Rep-1-Dpse_r2-4-modENCODE_2980-91 |
| exp11 | total-RNA-Developmental-Stage-Adult-Female-Whole-Species-Strain-D-pseudoobscura-wild-type-RNA-seq-Rep-1-Dpse_r2-4-modENCODE_3621-8 |
| exp12 | total-RNA-Developmental-Stage-Adult-Female-Whole-Species-Strain-D-pseudoobscura-wild-type-RNA-seq-Rep-1-Dpse_r2-4-modENCODE_3621-9 |
| exp13 | total-RNA-Developmental-Stage-Adult-Female-Whole-Species-Strain-D-pseudoobscura-wild-type-Tissue-Dpse-male-carcass-minus-reproductive-system-RNA-seq-Rep-1-Dpse_r2-4-modENCODE_4048-59 |
| exp14 | total-RNA-Developmental-Stage-Adult-Female-Whole-Species-Strain-D-pseudoobscura-wild-type-Tissue-Dpse-male-carcass-minus-reproductive-system-RNA-seq-Rep-1-Dpse_r2-4-modENCODE_4048-60 |
| exp15 | total-RNA-Developmental-Stage-Adult-Female-Whole-Species-Strain-D-pseudoobscura-wild-type-Tissue-Dpse-virgin-female-carcass-minus-reproductive-system-RNA-seq-Rep-1-Dpse_r2-4-modENCODE_4047-7 |
| exp16 | total-RNA-Developmental-Stage-Adult-Female-Whole-Species-Strain-D-pseudoobscura-wild-type-Tissue-Dpse-virgin-female-carcass-minus-reproductive-system-RNA-seq-Rep-1-Dpse_r2-4-modENCODE_4047-8 |
| exp17 | total-RNA-Developmental-Stage-Adult-Female-Whole-Species-Strain-D-pseudoobscura-wild-type-Tissue-Dpse-virgin-female-reproductive-system-RNA-seq-Rep-1-Dpse_r2-4-modENCODE_4049-1 |
| exp18 | total-RNA-Developmental-Stage-Adult-Female-Whole-Species-Strain-D-pseudoobscura-wild-type-Tissue-Dpse-virgin-female-reproductive-system-RNA-seq-Rep-1-Dpse_r2-4-modENCODE_4049-2 |
| exp19 | total-RNA-Developmental-Stage-Adult-Female-Whole-Species-Strain-D-pseudoobscura-wild-type-Tissue-Dpse-virgin-male-reproductive-system-RNA-seq-Rep-1-Dpse_r2-4-modENCODE_4050-3 |
| exp20 | total-RNA-Developmental-Stage-Adult-Female-Whole-Species-Strain-D-pseudoobscura-wild-type-Tissue-Dpse-virgin-male-reproductive-system-RNA-seq-Rep-1-Dpse_r2-4-modENCODE_4050-4 |
| exp21 | total-RNA-Developmental-Stage-Adult-Male-Strain-D-pseudoobscura-wild-type-Tissue-Dpse-Male-heads-RNA-seq-Rep-1-Dpse_r2-4-modENCODE_3085-792 |
| exp22 | total-RNA-Developmental-Stage-Adult-Male-Strain-D-pseudoobscura-wild-type-Tissue-Dpse-Male-heads-RNA-seq-Rep-1-Dpse_r2-4-modENCODE_3085-793 |
| exp23 | total-RNA-Developmental-Stage-Adult-Male-Strain-D-pseudoobscura-wild-type-Tissue-Dpse-Male-heads-RNA-seq-Rep-1-Dpse_r2-4-modENCODE_3085-794 |
| exp24 | total-RNA-Developmental-Stage-Adult-Male-Strain-D-pseudoobscura-wild-type-Tissue-Dpse-Male-heads-RNA-seq-Rep-1-Dpse_r2-4-modENCODE_3085-795 |
| exp25 | total-RNA-Developmental-Stage-Adult-Male-Strain-D-pseudoobscura-wild-type-Tissue-Dpse-Male-heads-RNA-seq-Rep-1-Dpse_r2-4-modENCODE_3085-796 |
| exp26 | total-RNA-Developmental-Stage-Adult-Male-Strain-D-pseudoobscura-wild-type-Tissue-Dpse-Male-heads-RNA-seq-Rep-1-Dpse_r2-4-modENCODE_3085-797 |
| exp27 | total-RNA-Developmental-Stage-Adult-Male-Strain-D-pseudoobscura-wild-type-Tissue-Dpse-Male-heads-RNA-seq-Rep-1-Dpse_r2-4-modENCODE_3085-798 |
| exp28 | total-RNA-Developmental-Stage-Adult-Male-Strain-D-pseudoobscura-wild-type-Tissue-Dpse-Male-heads-RNA-seq-Rep-1-Dpse_r2-4-modENCODE_3085-799 |
| exp29 | total-RNA-Developmental-Stage-Adult-Male-Strain-D-pseudoobscura-wild-type-Tissue-Dpse-Male-heads-RNA-seq-Rep-1-Dpse_r2-4-modENCODE_3085-800 |
| exp30 | total-RNA-Developmental-Stage-Adult-Male-Strain-D-pseudoobscura-wild-type-Tissue-Dpse-Male-heads-RNA-seq-Rep-1-Dpse_r2-4-modENCODE_3085-801 |
| exp31 | total-RNA-Developmental-Stage-Adult-Male-Whole-Species-Strain-D-pseudoobscura-wild-type-RNA-seq-Rep-1-Dpse_r2-4-modENCODE_3620-0 |
| exp32 | total-RNA-Developmental-Stage-Adult-Male-Whole-Species-Strain-D-pseudoobscura-wild-type-RNA-seq-Rep-1-Dpse_r2-4-modENCODE_3620-1 |

**Table S7.** The table shows the name of the datasets downloaded from modENCODE

| queryid | exp1 | exp2 | exp3 | exp4 | exp5 | exp6 | exp7 | exp8 | exp9 | exp10 | exp11 | exp12 | exp13 | exp14 | exp15 | exp16 |
|---|---|---|---|---|---|---|---|---|---|---|---|---|---|---|---|---|
| XP_002135331.1 | 0.00 | 0.00 | 0.00 | 8.35 | 0.00 | 0.00 | 3.58 | 0.00 | 0.00 | 0.00 | 0.0 | 0.0 | 0.00 | 0.0 | 0.0 | 0.00 |
| XP_015042266.1 | 19.57 | 11.90 | 27.36 | 18.10 | 27.46 | 18.00 | 20.69 | 10.90 | 27.40 | 20.90 | 0.0 | 0.0 | 0.00 | 0.0 | 0.0 | 0.00 |
| XP_002132370.1 | 2.17 | 0.00 | 0.00 | 0.00 | 0.00 | 0.00 | 0.00 | 0.00 | 0.00 | 0.00 | 0.0 | 0.0 | 0.00 | 0.0 | 0.0 | 0.00 |
| XP_015041143.1 | 1.53 | 2.09 | 1.92 | 0.00 | 2.90 | 1.06 | 2.73 | 0.77 | 0.00 | 0.00 | 0.0 | 0.0 | 0.00 | 0.0 | 0.0 | 0.00 |
| XP_002133104.2 | 3.08 | 0.00 | 23.22 | 0.00 | 0.00 | 0.00 | 14.63 | 9.25 | 4.84 | 3.28 | 0.0 | 0.0 | 0.00 | 0.0 | 0.0 | 0.00 |
| XP_002133789.1 | 4.71 | 6.44 | 5.93 | 6.54 | 0.00 | 0.00 | 5.60 | 0.00 | 2.47 | 2.52 | 0.0 | 0.0 | 0.00 | 0.0 | 0.0 | 0.00 |
| XP_015036675.1 | 21.00 | 9.14 | 15.62 | 9.27 | 19.29 | 13.17 | 12.49 | 19.15 | 16.79 | 13.00 | 0.0 | 0.0 | 0.00 | 0.0 | 0.0 | 0.00 |
| XP_015040408.1 | 789.76 | 540.08 | 745.24 | 730.52 | 387.83 | 635.71 | 495.66 | 956.97 | 915.80 | 849.21 | 0.0 | 0.0 | 0.00 | 0.0 | 0.0 | 350.08 |
| XP_015039586.1 | 0.00 | 0.00 | 0.00 | 0.00 | 0.00 | 0.00 | 0.00 | 0.00 | 0.00 | 0.00 | 0.0 | 0.0 | 0.00 | 0.0 | 0.0 | 0.00 |
| XP_015036957.1 | 98.72 | 110.47 | 135.50 | 99.62 | 79.33 | 74.30 | 106.72 | 63.00 | 70.69 | 74.27 | 0.0 | 0.0 | 178.22 | 0.0 | 0.0 | 0.00 |
| XP_002137838.2 | 0.00 | 0.00 | 0.00 | 0.00 | 0.00 | 0.00 | 0.00 | 0.00 | 0.00 | 0.00 | 0.0 | 0.0 | 0.00 | 0.0 | 0.0 | 0.00 |
| XP_015041911.1 | 7.33 | 11.45 | 7.90 | 5.81 | 2.64 | 2.89 | 4.98 | 8.40 | 9.89 | 6.71 | 0.0 | 0.0 | 0.00 | 0.0 | 0.0 | 0.00 |
| XP_015040782.1 | 4.82 | 8.80 | 0.00 | 8.92 | 4.06 | 13.31 | 0.00 | 4.84 | 5.91 | 1.72 | 0.0 | 0.0 | 0.00 | 0.0 | 0.0 | 0.00 |
| XP_015041370.1 | 2.95 | 4.04 | 0.00 | 0.00 | 0.00 | 0.00 | 3.51 | 2.96 | 2.32 | 2.36 | 0.0 | 0.0 | 0.00 | 0.0 | 0.0 | 0.00 |
| XP_015041786.1 | 0.00 | 0.00 | 0.00 | 0.00 | 0.00 | 0.00 | 0.00 | 0.00 | 3.53 | 3.59 | 0.0 | 0.0 | 0.00 | 0.0 | 0.0 | 0.00 |
| XP_015042146.1 | 2.58 | 0.00 | 6.50 | 14.34 | 9.79 | 10.70 | 0.00 | 11.66 | 8.14 | 12.42 | 0.0 | 0.0 | 0.00 | 0.0 | 0.0 | 0.00 |

| queryid | exp17 | exp18 | exp19 | exp20 | exp21 | exp22 | exp23 | exp24 | exp25 | exp26 | exp27 | exp28 | exp29 | exp30 | exp31 | exp32 |
|---|---|---|---|---|---|---|---|---|---|---|---|---|---|---|---|---|
| XP_002135331.1 | 0.00 | 0.0 | 0.0 | 0.00 | 1.38 | 0.00 | 0.00 | 0.00 | 0.00 | 0.00 | 0.00 | 0.00 | 0.63 | 0.00 | 0.0 | 0.0 |
| XP_015042266.1 | 0.00 | 0.0 | 0.0 | 0.00 | 9.99 | 20.42 | 22.24 | 8.62 | 35.05 | 16.26 | 16.23 | 22.41 | 25.64 | 18.46 | 0.0 | 0.0 |
| XP_002132370.1 | 0.00 | 0.0 | 0.0 | 0.00 | 0.00 | 0.00 | 0.00 | 0.00 | 0.00 | 0.00 | 0.00 | 0.00 | 0.00 | 0.00 | 0.0 | 0.0 |
| XP_015041143.1 | 0.00 | 0.0 | 0.0 | 0.00 | 2.11 | 0.51 | 1.12 | 1.52 | 0.00 | 0.95 | 3.81 | 0.36 | 0.32 | 0.18 | 0.0 | 0.0 |
| XP_002133104.2 | 0.00 | 0.0 | 0.0 | 0.00 | 0.00 | 0.00 | 4.49 | 0.00 | 0.00 | 7.67 | 7.65 | 0.00 | 5.18 | 5.80 | 0.0 | 0.0 |
| XP_002133789.1 | 0.00 | 0.0 | 0.0 | 0.00 | 4.33 | 3.16 | 0.00 | 0.00 | 0.00 | 0.00 | 0.00 | 3.31 | 0.00 | 2.22 | 0.0 | 0.0 |
| XP_015036675.1 | 0.00 | 0.0 | 0.0 | 0.00 | 10.53 | 5.76 | 8.37 | 9.46 | 3.85 | 8.33 | 9.50 | 11.41 | 9.05 | 7.65 | 0.0 | 0.0 |
| XP_015040408.1 | 553.07 | 0.0 | 0.0 | 495.78 | 715.48 | 632.77 | 577.04 | 304.33 | 441.99 | 519.54 | 409.46 | 919.94 | 706.68 | 812.05 | 0.0 | 0.0 |
| XP_015039586.1 | 0.00 | 0.0 | 0.0 | 0.00 | 0.00 | 0.00 | 0.00 | 4.60 | 0.00 | 0.00 | 0.00 | 0.00 | 0.00 | 0.00 | 0.0 | 0.0 |
| XP_015036957.1 | 0.00 | 0.0 | 0.0 | 0.00 | 169.02 | 162.54 | 203.27 | 302.36 | 307.38 | 156.61 | 245.68 | 163.99 | 170.06 | 154.46 | 0.0 | 0.0 |
| XP_002137838.2 | 0.00 | 0.0 | 0.0 | 0.00 | 0.00 | 0.00 | 0.00 | 0.00 | 0.00 | 0.00 | 0.00 | 0.00 | 0.00 | 1.54 | 0.0 | 0.0 |
| XP_015041911.1 | 0.00 | 0.0 | 0.0 | 0.00 | 7.69 | 7.02 | 10.71 | 0.00 | 8.44 | 5.22 | 7.82 | 9.81 | 11.02 | 8.89 | 0.0 | 0.0 |
| XP_015040782.1 | 0.00 | 0.0 | 0.0 | 0.00 | 2.95 | 10.79 | 2.35 | 12.75 | 12.96 | 0.00 | 4.00 | 3.77 | 2.71 | 3.79 | 0.0 | 0.0 |
| XP_015041370.1 | 0.00 | 0.0 | 0.0 | 0.00 | 1.36 | 3.96 | 4.31 | 0.00 | 5.95 | 0.00 | 0.00 | 4.15 | 4.35 | 4.17 | 0.0 | 0.0 |
| XP_015041786.1 | 0.00 | 0.0 | 0.0 | 0.00 | 0.00 | 0.00 | 0.00 | 0.00 | 0.00 | 0.00 | 0.00 | 3.15 | 0.00 | 6.33 | 0.0 | 0.0 |
| XP_015042146.1 | 0.00 | 0.0 | 0.0 | 0.00 | 9.50 | 6.93 | 11.33 | 5.12 | 0.00 | 3.22 | 9.65 | 9.69 | 8.71 | 11.58 | 0.0 | 0.0 |

**Table S8.** For the 16 *D. pseudoobscura* genes out of the 38 orphans found by blastp+tblastn, it is shown the expression level in 32 different RNA-Seq datasets. The values in the columns corresponds to the RNA expression data in Reads Per Kilobase Million (RPKM).

| Method | Gene ID | Taxon | Hit E-value |
|---|---|---|---|
| BLASTP (17/54) | | | |
| | NP_608541.1 | Metarhizium robertsii | 0.00 |
| | NP_611823.2 | Acinetobacter guillouiae | $7.59 \times 10^{-6}$ |
| | NP_001014551.1 | Daphnia pulex | $8.89 \times 10^{-5}$ |
| | NP_652358.1 | Drosophila simulans | $4.17 \times 10^{-7}$ |
| | NP_001097701.1 | Drosophila simulans | $9.85 \times 10^{-20}$ |
| | NP_001245834.1 | Drosophila simulans | $8.78 \times 10^{-14}$ |
| | NP_001261798.1 | Drosophila simulans | $3.33 \times 10^{-6}$ |
| | NP_001163438.1 | Drosophila simulans | $3.33 \times 10^{-6}$ |
| | NP_001163495.1 | Drosophila simulans | $1.02 \times 10^{-10}$ |
| | NP_001162981.1 | Drosophila simulans | $6.39 \times 10^{-11}$ |
| | NP_001189120.1 | Drosophila simulans | 0.00 |
| | NP_001259862.1 | Drosophila simulans | $8.78 \times 10^{-14}$ |
| | NP_001097010.1 | Drosophila simulans | $8.08 \times 10^{-103}$ |
| | NP_001286780.1 | Acinetobacter guillouiae | $7.59 \times 10^{-6}$ |
| | NP_001097828.1 | Drosophila ananassae | $3.85 \times 10^{-6}$ |
| | NP_001259864.1 | Drosophila simulans | $4.17 \times 10^{-7}$ |
| | NP_524254.1 | Drosophila simulans | $9.85 \times 10^{-20}$ |

**Table S9.** 54 genes deemed as orphans in *Drosophila melanogaster* by blastp+tblastn, are searched in GenBank NonRedundant (nr) database, using two BLAST variants: blastp and tblastn. For each of these searches, it is reported the number (in parentheses) of queries with at least one hit with E-value < 0.001. For each gene, it is shown the E-value of its best hit in nr, and the target species.

| Method | Gene ID | Taxon | Hit E-value |
|---|---|---|---|
| BLASTP (2/38) | | | |
| | XP_015039586.1 | Drosophila grimshawi | 0.00 |
| | XP_015041911.1 | Patagioenas fasciata monilis | 0.00 |
| TBLASTN (1/38) | | | |
| | XP_015039586.1 | Drosophila melanogaster | $1.06 \times 10^{-6}$ |

**Table S10.** 38 genes deemed as orphans in *Drosophila pseudoobscura* by blastp+tblastn, are searched in GenBank NonRedundant (nr) database, using two BLAST variants: blastp and tblastn. For each of these searches, it is reported the number (in parentheses) of queries with at least one hit with E-value < 0.001. For each gene, it is shown the E-value of its best hit in nr, and the target species.

| E-value | blastp | tblastn |
|---|---|---|
| 0.10000 | 248 | 132 |
| 0.01000 | 24 | 15 |
| 0.00100 | 2 | 1 |
| 0.00010 | 0 | 0 |
| 0.00001 | 0 | 0 |

**Table S11.** False positive orphans found but blastp and tblastn at different E-values thresholds. These are the hits obtained using as queries scrambled sequences of the original ones.
